# YAP/TEAD drives treatment-induced adaptive immunosuppression in EGFR-mutant lung cancer

**DOI:** 10.64898/2026.01.22.701073

**Authors:** Meija Honkanen, Jonas Baumgarten, Nikol Dibus, Jenna H. Rannikko, Milos Gojkovic, Paloma Cejas, Yingtian Xie, Zsuzsanna Ortutay, Johannes Merilahti, Mari Tienhaara, Iman Farahani, Steffen Boettcher, Pasi A. Jänne, Otto Kauko, Raphael A. Nemenoff, Lynn E. Heasley, Heidi M. Haikala, Maija Hollmén, Kari J. Kurppa

## Abstract

Residual disease remains a major obstacle for achieving durable responses in patients treated with oncogene-targeted therapy. Drug-tolerant persister (DTP) cells emerging under treatment and persisting in residual tumors are considered to be the root of acquired resistance, yet their contribution to immune evasion in on-treatment tumors is poorly defined. Here, we show in the context of EGFR-mutant lung cancer that DTP cells actively contribute to the formation of an immunosuppressive tumor microenvironment during EGFR tyrosine kinase inhibitor (TKI) therapy. In syngeneic mouse models and in patients, EGFR TKI therapy leads to an accumulation of immunosuppressive macrophages, which is strictly treatment-dependent and fully reversible upon treatment cessation or progressive disease, respectively. Quiescent DTP cells directly drive the recruitment and immunosuppressive reprogramming of monocytes and macrophages through a YAP-driven secretome, and the DTP-reprogrammed monocytes suppress T cell proliferation and effector functions *in vitro*. Co-targeting YAP with a TEAD inhibitor ORM-47286 rewires the DTP secretome and inhibits macrophage reprogramming *in vitro*, and prevents immunosuppressive macrophage accumulation and improves the efficacy of EGFR TKI therapy in immunocompetent mouse models. Our findings highlight the previously unappreciated role of DTP cells in modulating the tumor microenvironment in on-treatment tumors, and position the treatment-induced YAP/TEAD activity in DTP cells as an important driver of adaptive immunosuppression during EGFR-targeted therapy.

## Introduction

During initial response to oncogene-targeted therapy, cancer cells can undergo non-genetic adaptation that allows them to remain quiescent in residual tumors. These drug-tolerant persister (DTP) cells serve as seeds for drug resistance, and can acquire additional mechanisms of resistance over time, ultimately leading to treatment relapse.^1,2^ It has become evident that drug-tolerant cells are phenotypically distinct from proliferating, untreated tumor cells. Accumulating evidence across different oncogene-addicted cancer types suggests that the acquired DTP cell states are associated with cancer cell-intrinsic inflammatory response, a reversible senescence-like phenotype, and increased expression of genes encoding secreted factors.^3–6^ These changes, many of which are linked to tumor–microenvironment crosstalk, raise the question whether on-treatment cancer cells can shape the residual tumor microenvironment and, especially, actively modulate the immune response. However, current insights into drug tolerance come mostly from *in vitro* studies and xenograft models that limit our understanding of the role of tumor immune microenvironment in this process.

Treatment-induced activation of the transcriptional co-activator Yes-associated protein 1 (YAP) has emerged as a major mediator of resistance to oncogene-targeted therapy, particularly in EGFR- and KRAS-driven lung cancers.^7–9^ In EGFR-mutant lung cancer, YAP activation has been shown to drive cell-intrinsic mechanisms that suppress the initial drug-induced apoptosis, facilitating the establishment of the DTP state.^7^ Preventing treatment-induced YAP activation with upfront combination therapies using TEAD inhibitors markedly reduces the number of surviving DTP cells and improves the efficacy of EGFR-targeted therapy.^7,9^ Intriguingly, YAP also has extensive non–cell-autonomous functions.^10^ In particular, YAP activation has been shown to drive the secretion of cytokines and chemokines that suppress antitumor immunity by recruiting tumor-associated macrophages to the tumor microenvironment.^11–15^ Hence, treatment-induced YAP activity in DTP cells could also promote microenvironmental mechanisms of resistance.

Here, we investigated how the DTP cells contribute to the remodeling of the tumor immune microenvironment in EGFR-mutant lung cancer. We show that on-treatment, quiescent DTP cells actively orchestrate immunosuppressive macrophage reprogramming through YAP-driven expression of secreted factors, and that the DTP-educated monocytes suppress T-cell activation. Blocking treatment-induced YAP activity with upfront TEAD inhibitor combination therapy *in vivo* prevented EGFR-targeted therapy–induced recruitment of immunosuppressive macrophages, and improved the efficacy of EGFR-targeted therapy in syngeneic mouse models. Together, our results suggest that DTP cells are active modulators of the tumor microenvironment in residual tumors and can directly manipulate the tumor immune response to promote immunosuppression.

## Results

### EGFR-targeted therapy leads to the accumulation of immunosuppressive macrophages in residual tumors

In order to assess treatment-induced changes within the tumor microenvironment of residual tumors following EGFR-targeted therapy, we utilized two syngeneic mouse models of EGFR mutant lung cancer, del19.2 and L860R^16^ – representing the two most common EGFR activating mutations observed in patients; exon 19 deletion and L858R mutations, respectively. To this end, we treated tumor-bearing mice for three weeks with EGFR TKI osimertinib (**Figure 1A**). The treatment resulted in dramatic tumor regression in both models, with nearly impalpable tumors after three weeks of treatment. However, viable residual DTP cells persisted in both models despite sustained treatment response, as evidenced by the fast regrowth of tumors upon treatment cessation (**Figure 1A**). Immunohistochemical analysis revealed an immunologically “cold” phenotype with sparse T cell infiltration in vehicle-treated tumors, typical for non-small cell lung cancer (NSCLC) tumors harboring EGFR driver mutations.^17^ In contrast, we observed a significant treatment-induced infiltration of cytotoxic CD8^+^ T cells in the residual tumors in both models (**Figure 1B, C**), demonstrating that the on-treatment, residual tumors are immunogenic and stimulate a T cell response. However, the CD8^+^ T cell response was accompanied by a massive accumulation of F4/80^+^ macrophages in the tumor microenvironment. These macrophages expressed markers of immunosuppressive M2-like phenotype, CD206 and/or ARG-1 (**Figure 1B, C**). Strikingly, following treatment cessation and tumor regrowth, both macrophage and T cell numbers returned to the levels observed in vehicle-treated tumors, suggesting that both T cell and macrophage recruitment are highly specific to the residual, on-treatment state (**Figure 1B, C**). We were also able to recapitulate the treatment-specific recruitment of immunosuppressive macrophages in a PC-9 xenograft model using nude mice, demonstrating that this phenomenon is not dependent on the T cell response (**Figure 1D-F**). Furthermore, we observed that the treatment-induced recruitment of immunosuppressive macrophages was dose-dependent – a daily 10 mg/kg osimertinib dose that resulted in a deep and sustained response promoted more robust accumulation of F4/80^+^ and CD206^+^ macrophages to the residual tumors than a suboptimal 2.5 mg/kg dose that resulted in poor efficacy and transient tumor control (**Figure 1D-F**). This result is consistent with our observations in the syngeneic models and emphasizes that the recruitment of immunosuppressive macrophages strongly associates with a non-proliferative, residual tumor state. Together, these observations demonstrate that EGFR-targeted therapy promotes the induction of an immunosuppressive microenvironment in the residual tumors, highlighted by the recruitment of immunosuppressive macrophages, and that these changes are specific to a quiescent, on-treatment state.

**Figure 1.**
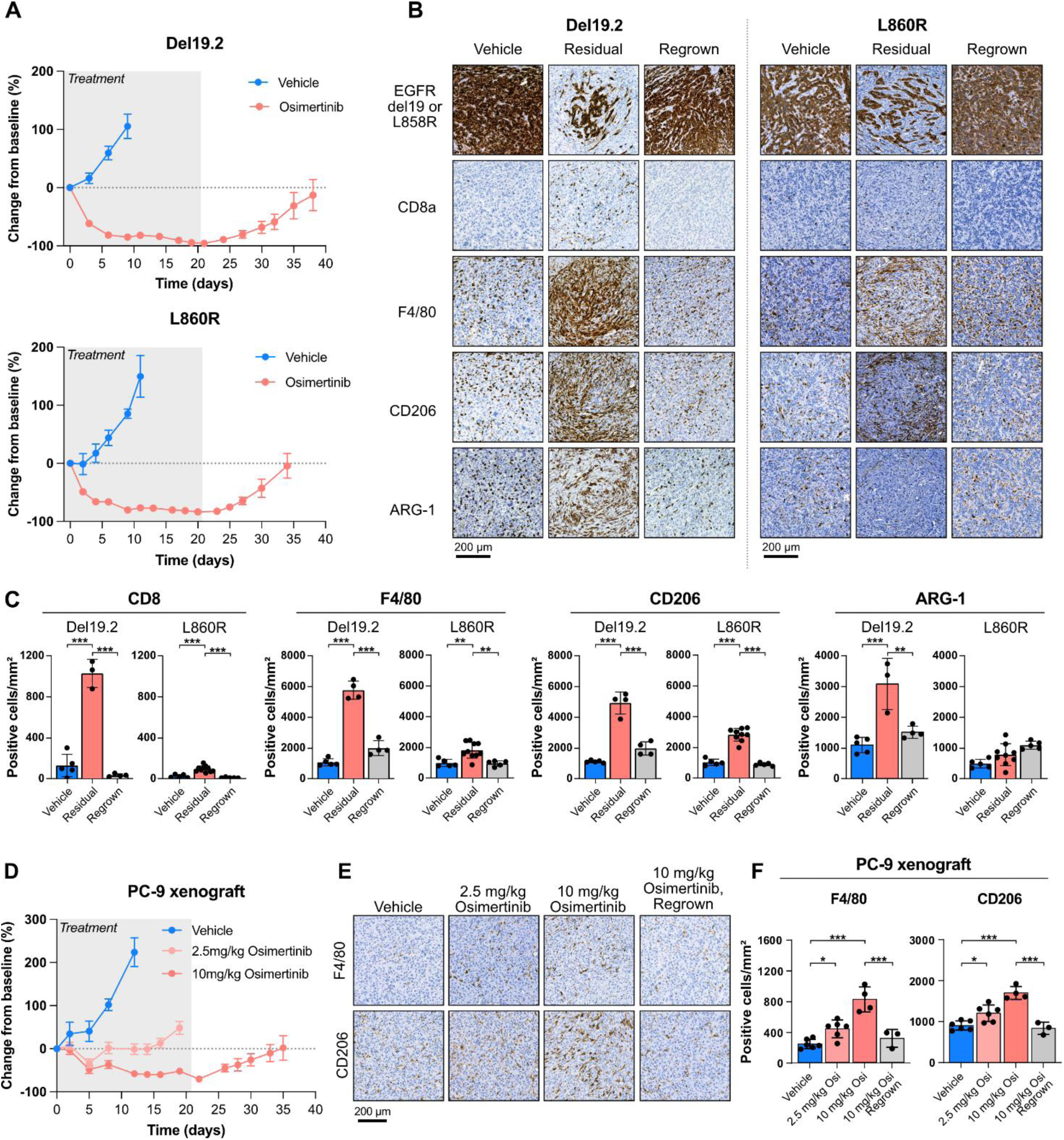
EGFR-targeted therapy leads to infiltration of T-cells and immunosuppressive macrophages into the tumor microenvironment in syngeneic mouse models of EGFR-mutant NSCLC. A) Osimertinib response in syngeneic EGFR-mutant lung cancer models Del19.2 and L860R in C57Bl/6 mice (n = 10 mice). B) Immunohistochemical staining of tumor samples from (A). EGFRdel19, cancer cells; CD8a, cytotoxic T-cells; F4/80, macrophages; CD206, M2 macrophage marker. C) Quantification of positive cells detected per mm^2^ in tumors after 3-weeks of treatment depicted in (B) using QuPath. D) Osimertinib response in PC-9 xenograft model in nude mice. E) Immunohistochemical staining of (D). Quantification of (E) performed as in (C). Data shown as mean ± SEM (A, D) or mean ± SD (C, F). One-way ANOVA was used to assess statistical significance. ***, P-value <0.001; **, P-value <0.01; *, P-value < 0.05.

### Secreted factors from quiescent, on-treatment cancer cells recruit and reprogram monocytes and macrophages to an immunosuppressive state

Quiescent, residual DTP cells have been shown to adopt a reversible senescence-like phenotype, characterized by highly increased secretion of cytokines and chemokines.^3,4,7^ We hypothesized that this secretion of immunomodulatory factors could directly drive the immunosuppressive polarization and/or recruitment of monocytes/macrophages we observed *in vivo*. To test this hypothesis, we exposed THP-1 monocytes to media conditioned by DTP cells or by proliferating, untreated control cells (**Figure 2A**). Since the accumulation of immunosuppressive macrophages *in vivo* was associated with sustained tumor growth control (**Figure 1D-E**), we sought to model the corresponding quiescent state *in vitro*. To this extent, we established DTP cells by treating EGFR-mutant NSCLC cells with either single-agent osimertinib, or with a combination of osimertinib and MEK inhibitor trametinib. Single-agent osimertinib-treated cells quickly re-activate the MAPK pathway *in vitro* and resume cell cycle (albeit at a low rate), whereas the combined EGFR/MEK inhibition maintains a stable, quiescent drug-tolerant state over a long period of time (**Figure 2B, Supplementary Figure S1A**).^7,18,19^ We observed a robust increase in the expression of immunosuppressive macrophage markers *CD163* and *MRC1* (CD206) in THP-1 cells exposed to conditioned medium from osimertinib/trametinib-treated cells, compared to medium from proliferating, DMSO-treated cells (**Figure 2C, Supplementary Figure S1B**). However, the induction of these markers in THP-1 cells exposed to conditioned medium from single-agent osimertinib-treated cells was significantly lower (**Figure 2C, Supplementary Figure S1B**), consistent with our *in vivo* observation that this phenomenon is specific to a quiescent residual state. Indeed, when the osimertinib/trametinib-treated DTP cells (hereafter “DTP cells”) underwent either a short (48 hours) or long (13 days) drug holiday, the ability of the conditioned medium from these cells to promote the expression of *CD163* or *MRC1* in THP-1 cells was drastically reduced (**Figure 2D**). Importantly, the drugs alone did not affect the expression of these markers in THP-1 cells (**Supplementary Figure S1C**). We also assessed whether secreted factors from DTP cells can promote the recruitment of monocytes by performing chemotaxis assays using human CD14^+^ peripheral blood monocytes from healthy donors. The monocytes migrated significantly more towards conditioned medium from osimertinib/trametinib-treated DTP cells than conditioned medium from DMSO-treated cells, demonstrating that the DTP cells have an enhanced ability to recruit monocytes (**Figure 2E)**. Analysis of a broad cytokine and chemokine panel revealed a conserved upregulation of immunomodulatory and monocyte-recruiting cues across three EGFR-mutant NSCLC lines in the DTP state (**Supplementary Figure S1D, E).** This secretory profile aligns with the enhanced macrophage accumulation and immunosuppressive bias we observed *in vivo* (**Figure 1B-C**). Together, these results supported our hypothesis and suggested that secreted factors from DTP cells can promote the recruitment of monocytes and their subsequent immunosuppressive polarization, but this ability is dependent on a sustained quiescent state.

**Figure 2.**
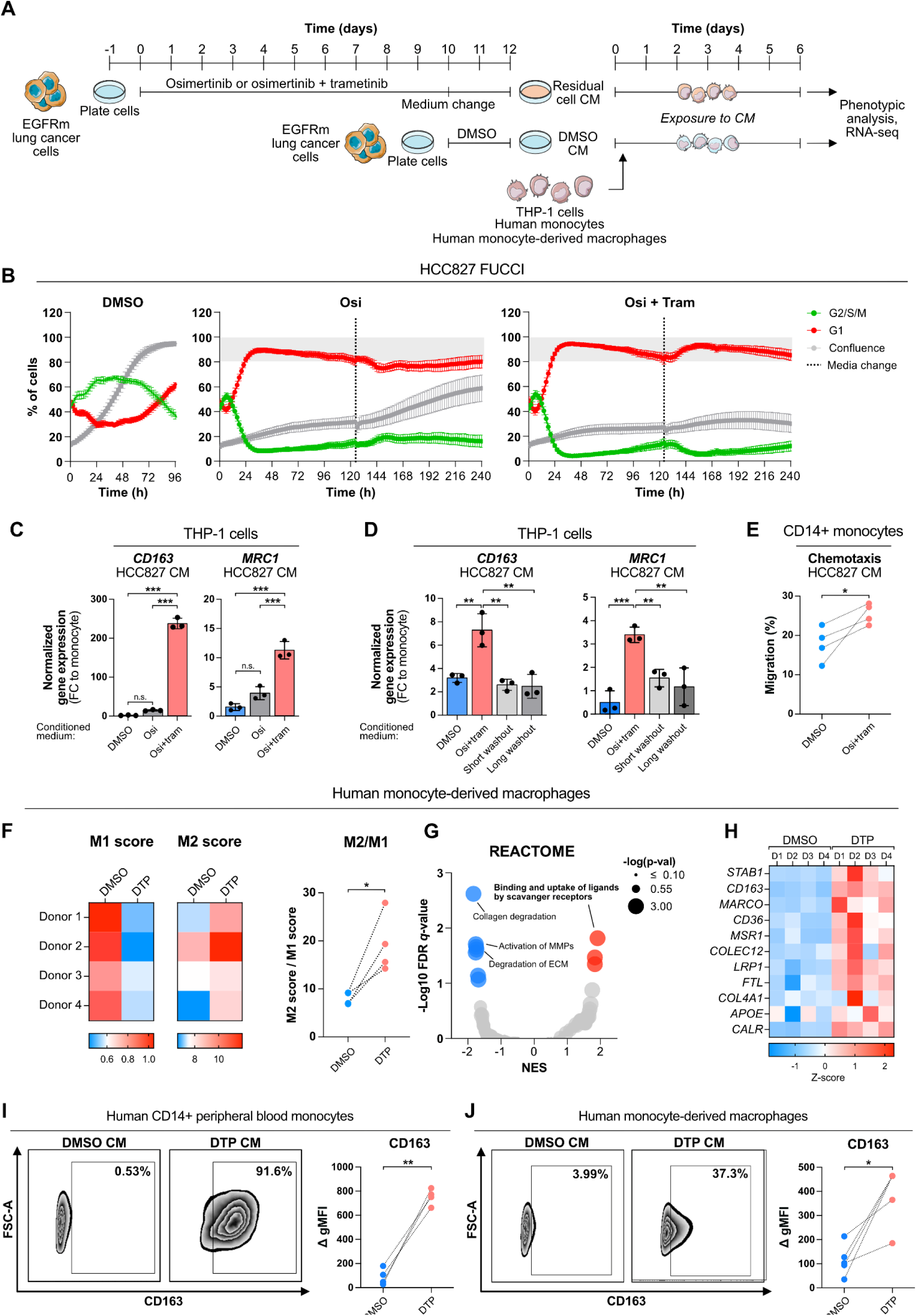
Secreted factors from quiescent, on-treatment cancer cells reprogram macrophages to an M2-like immunosuppressive state. A) Experimental set-up to generate DTP cells, collection of CM, and exposure to monocytes and macrophages. B) HCC827 cells expressing pLEX307-FUCCI cell cycle reporter were treated with either DMSO, 100 nM osimertinib alone, or in combination with 30 nM trametinib for 10 days. C) THP-1 cells were treated as in (B), and the expression of M2 marker genes (CD163 and MRC1/CD206) was analyzed with RT-qPCR. D) CM was collected as in (B), with the addition of either short (48 h) or long (13 days) drug holidays, followed by THP-1 incubation as in (C). E) CD14⁺ monocyte chemotaxis toward CM from DMSO- or osimertinib/trametinib–treated cancer cells. Chemotaxis was measured using Incucyte live-cell imaging. Data shown as a percentage of cells migrated after 24 hours. F) RNA-sequencing of monocyte-derived macrophages treated according to (A), with M1 and M2 gene set^21^ expression scored relative to expression of all genes. G) GSEA of REACTOME pathways in monocyte-derived macrophages treated as in (F). H) Relative expression of genes driving the enrichment of binding and uptake of ligands by scavenger receptors -gene set (G). I) Flow cytometry analysis of cell surface expression of CD163 in monocytes and monocyte-derived macrophages (J) exposed to DTP CM. DTP = drug-tolerant persister cells, achieved with osimertinib/trametinib treatment for 12 days, CM = conditioned medium, GSEA = gene set enrichment analysis. Data shown as mean ± SD in C and D. One-way ANOVA (C, D) or paired t-test (E, F, I, J) was used to assess statistical significance. ***, P-value <0.001; **, P-value <0.01; *, P-value < 0.05

Our results suggest that as monocytes recruited to the on-treatment tumor differentiate to macrophages, they encounter a niche rich in secreted factors from quiescent DTP cells. Therefore, we wanted to assess if the DTP cells can promote phenotypic changes also in differentiated macrophages. For this, we analyzed the transcriptomic changes promoted by the conditioned medium from osimertinib/trametinib-treated DTP cells in human monocyte-derived macrophages using RNA-sequencing. Consistent with our results in THP-1 monocytes, the exposure to DTP cell conditioned medium promoted a transcriptomic reprogramming of macrophages into an immunosuppressive, M2-like state (**Figure 2F**). Furthermore, a GSEA analysis of the REACTOME pathway database^20^ indicated that genes involved in the binding and uptake of ligands by scavenger receptors were highly enriched in macrophages exposed to the DTP cell-conditioned medium. A more detailed analysis of the genes primarily driving the enrichment of this pathway revealed a significantly increased expression of several macrophage scavenger receptor genes associated with immunosuppressive macrophages, including *CD163*, *STAB1* (CLEVER-1), and *MARCO* (**Figure 2G, H**). Indeed, consistent with the upregulation of *CD163* in THP-1 cells, the exposure to the residual cancer cell medium promoted a robust increase in CD163 surface expression in both human monocytes and monocyte-derived macrophages (**Figure 2I, J**).

Together, these data demonstrate that the quiescent, residual DTP cells can actively recruit monocytes and drive the immunosuppressive phenotypic reprogramming of monocytes/macrophages through the increased secretion of soluble factors.

### DTP-reprogrammed monocytes suppress T cell activation and proliferation

To assess the functional consequence of DTP-mediated reprogramming of monocytes/macrophages on adaptive immune responses, we investigated how DTP cell secretome-educated monocytes influence T cell activation *in vitro* (**Figure 3A**). We isolated monocytes from healthy blood donors and exposed them to DTP-conditioned media derived from HCC827 cells for six days. After exposure, monocytes were co-cultured for 72 hours with pre-activated T cells from matched donors and analyzed for proliferation and activation by flow cytometry. We observed that the DTP-reprogrammed monocytes suppressed the proliferation of CD4^+^ and CD8^+^ T cells and promoted a consistent decrease in the expression of markers for T cell activation; IFN-ɣ and granzyme B in CD8^+^ T cells, and IFN-ɣ and TNF in CD4^+^ T cells. (**Figure 3B, C, Supplementary Figure S2**). These data demonstrate that monocytes reprogrammed by DTP secretome acquire the capacity to dampen T cell proliferation and effector functions, potentially to support immune evasion in residual tumors.

**Figure 3.**
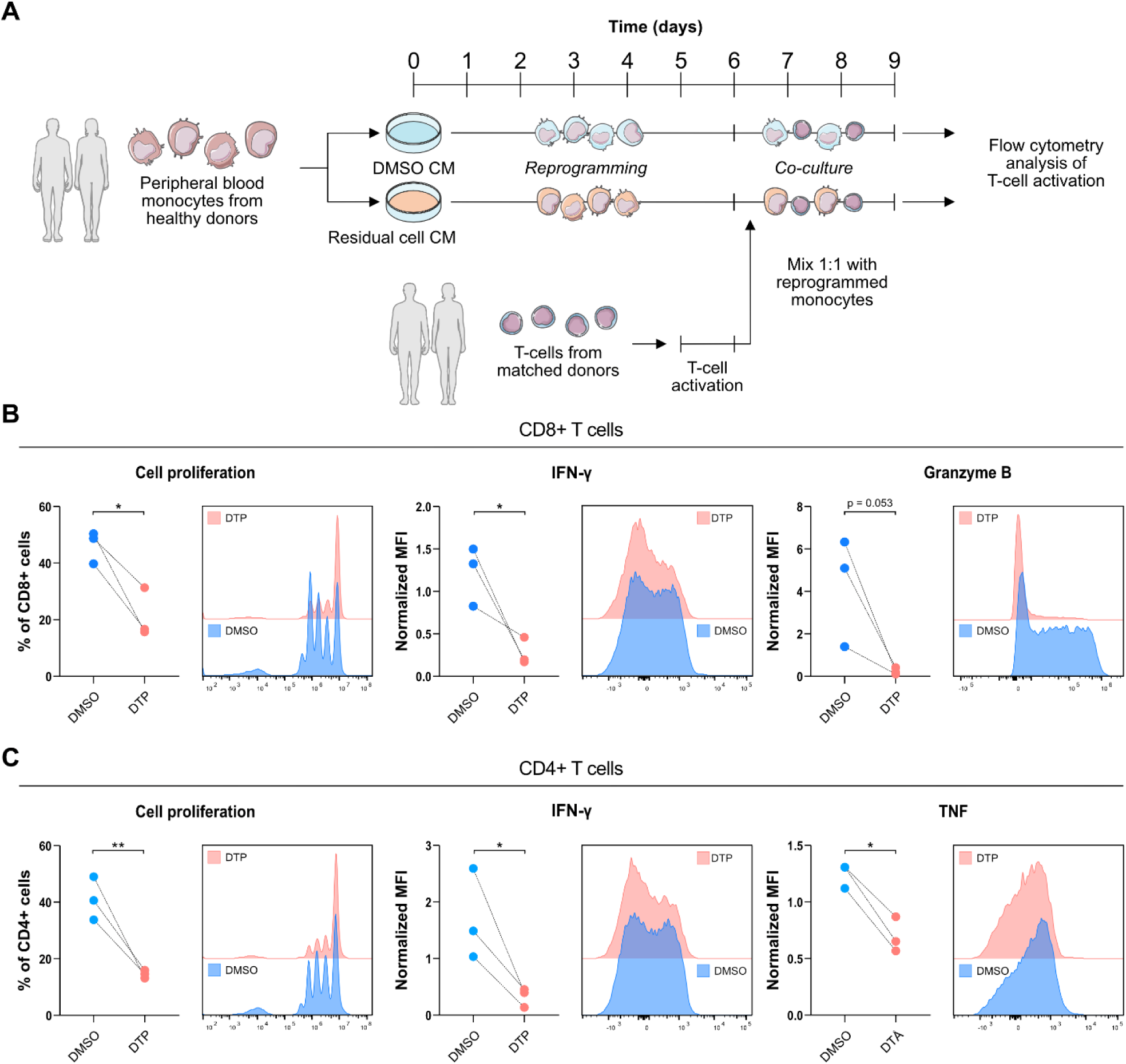
DTP-reprogrammed monocytes suppress T cell activation and proliferation. A) Experimental set-up for monocyte and T cell co-culture experiments. Monocytes were isolated from healthy blood donors and exposed to HCC827 DTP or control CM. After 6 days of exposure to CM, monocytes were co-cultured with activated CD3^+^ T cells isolated from the same donors for 72 hours, followed by analysis of CD8+ (B) and CD4^+^ (C) T cell proliferation and activation by flow cytometry. CM = conditioned medium. A t-test was used to assess statistical significance. **, P-value <0.01; *, P-value < 0.05

### CSF1-R -inhibitor pexidartinib selectively depletes proinflammatory macrophages and suppresses T cell infiltration *in vivo* when combined with osimertinib

Since our data suggested that macrophages recruited to on-treatment tumors may contribute to immunosuppression, we next explored *in vivo* whether depleting macrophages could enhance the efficacy of EGFR inhibitor treatment. Given the established role of CSF1-CSF1R signaling in n regulating tumor-associated macrophage recruitment^22^, we tested the CSF1-R inhibitor pexidartinib^23^ in combination with osimertinib *in vivo*. Unexpectedly, the combination of pexidartinib and osimertinib did not result in therapeutic benefit over osimertinib monotherapy in the L860R model, the combination being in fact marginally worse than osimertinib alone (**Supplementary Figure S3A, B**). Immunohistochemical analysis clearly demonstrated that co-administered pexidartinib was able to reduce the number of F4/80^+^ tumor-associated macrophages (TAM) in the osimertinib-treated tumors, but surprisingly also led to a significant decrease in number of infiltrating CD8^+^ T-cells (**Supplementary Figure S3C, D**). More detailed flow cytometry -based immune profiling of tumors harvested after four days of treatment demonstrated that while pexidartinib indeed effectively reduced the number of CD11b^+^/Ly6G^-^ TAMs, this effect was driven by the specific depletion of pro-inflammatory, M1-like CD11b^+^/Ly6G^-^/MHCII^high^ macrophages, leaving the immunosuppressive, M2-like CD11b^+^/Ly6G^-^/MHCII^low^ macrophages unaffected (**Supplementary Figure S3E, F**). This selective depletion of M1-like macrophages led to a significantly increased M2/M1 macrophage ratio compared to osimertinib monotherapy (**Supplementary Figure S3E**), and may explain the reduced infiltration of T cells in pexidartinib + osimertinib-treated tumors, also evident in the flow cytometry analysis as a decrease in CD45^+^/CD3^+^ cells (**Supplementary Figure S3E**).

### Immunosuppressive macrophages are enriched in on-treatment tumors of EGFR-mutant NSCLC patients treated with EGFR inhibitors

Our results from *in vivo* and *in vitro* models suggested a DTP-driven, and residual tumor-specific recruitment and reprogramming of monocytes and macrophages (MoMacs) in on-treatment tumors. To assess if similar phenotypic changes also occur in the tumor immune microenvironment in patients, we used a published single-cell RNA-sequencing dataset of longitudinally sampled (treatment naïve, TN; residual disease, RD; progressive disease, PD) lung cancer patients who underwent oncogene-targeted therapy.^24^ After pre-processing, main cell type clustering and immune cell subclustering (**Supplementary Figure S4**, **Supplementary Table S1**, and materials and methods), we focused exclusively on MoMacs in samples from EGFR-mutant lung cancer patients (n=16 patients, 26 samples), and further curated the samples by including only 1) samples with sufficient (>15) MoMacs for analysis (n=11 patients, 17 samples), and 2) lung tumor samples (n=6 patients, 9 samples). Of these nine samples, three were from TN tumors, five from RD, and one from PD. All included patients were treated with osimertinib.

Initial clustering of MoMacs from all nine samples (n = 604 cells) identified six distinct clusters (**Figure 4A**), with the MoMacs from TN tumors and from the PD sample enriching in cluster 0, and MoMacs from RD tumors enriching mainly in clusters 1, 2, and 3 (**Figure 4B, C**). The clustering was not driven by patient, biopsy type, or experimental batch (**Figure 4C**). The enrichment of TN, RD, and PD MoMacs in different clusters suggested that the MoMacs in RD samples were transcriptionally, and thus potentially phenotypically, different from those in TN and PD samples, supporting our *in vitro* and *in vivo* findings. In addition, while our curated sample set contained only two patients with samples from more than one timepoint (TN-RD-PD; TN-RD), in both cases the majority of TN MoMacs were enriched in cluster 0, while RD MoMacs mainly resided in clusters 1, 2, and 3 (**Figure 4B**). In the case of one patient who had samples from all timepoints, the MoMacs in the PD sample were enriched in cluster 0 together with the TN MoMacs (**Figure 4B**), suggesting that the enrichment of TN, RD, and PD MoMacs in distinct clusters is not driven by MoMac heterogeneity in different patient samples, but is a consequence of response to EGFR-targeted therapy.

**Figure 4.**
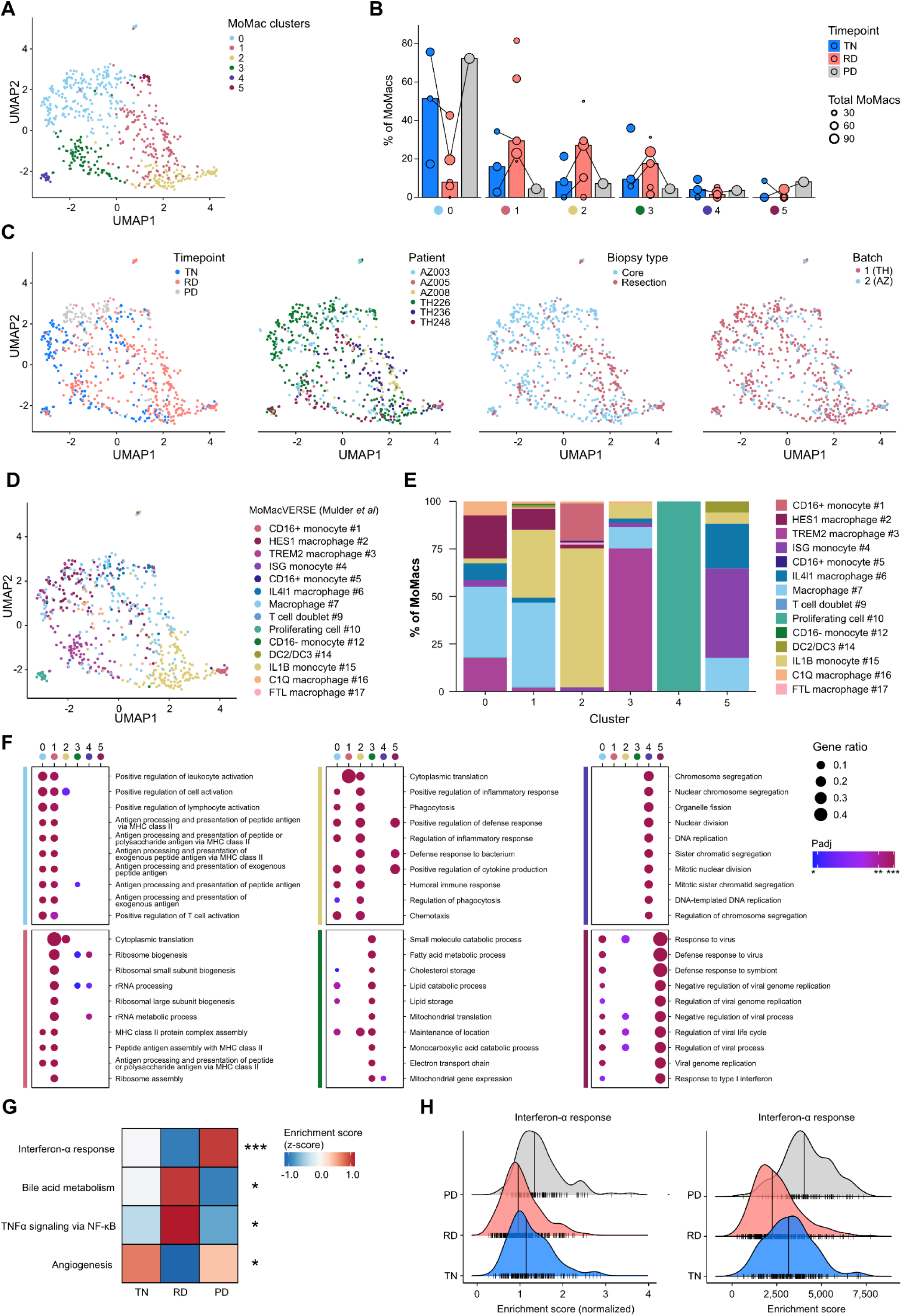
Immunosuppressive macrophages are enriched in on-treatment tumors of EGFR-mutant NSCLC patients treated with EGFR inhibitors. A) UMAP plot showing clustering of MoMacs (n = 604 cells, n = 9 biopsies) from lung biopsies of lung adenocarcinoma patients with EGRF driver gene. B) Bars show proportion of cells in each MoMac cluster by targeted therapy status (median). Dots represent individual patient biopsies (TN: n = 3; RD: n = 5; PD: n = 1) with diameter relative to total MoMac number in biopsy. Lines connect biopsies from the same patient. C) UMAP plot from A colored by patient, targeted therapy status, experimental batch or biopsy type. D) MoMacs were mapped to a human monocyte and macrophage single-cell atlas MoMacVERSE^25^ and original UMAP plot from A was colored by predicted MoMacVERSE cell types. E) Proportion of cells in UMAP clusters from A mapping to each MoMacVERSE cell type. F) GO biological process enrichment analysis on DEGs of each MoMac cluster. Dot plots show top 10 significantly enriched (Padj and q-value <0.05) biological processes in each MoMac cluster. G and H) Hallmark gene set enrichment analysis. A heatmap of normalized enrichment scores of gene sets significantly differing between TN and RD (G). Asterisks indicate adjusted P-values for Wilcoxon rank sum test between TN and RD group. Ridge plots showing distribution of normalized (left) and unnormalized (right) enrichment scores in TN, RD and PD group (H). Vertical lines indicate group median and vertical ticks display scores for individual cells. PD, progressive disease; RD, residual disease; TN, targeted therapy naïve; UMAP, uniform manifold approximation and projection. ***, P-value <0.001; *, P-value <0.05.

To understand the phenotypic differences underlying the differential clustering of TN and RD MoMacs, we utilized the MoMacVERSE single-cell atlas of human monocytes and macrophages^25^, and annotated each cell with a MoMacVERSE subtype based on transcriptional similarity (**Figure 4D** and **Supplementary Figure S5A**). Intriguingly, while the TN-enriched cluster 0 harbored a relatively heterogenous population of monocytes and macrophages, the RD-enriched cluster 1, and especially cluster 2, were overrepresented by cells with IL1B and CD16^+^ monocyte signatures (**Figure 4D, E**), potentially reflecting increased recruitment of monocytes to the on-treatment tumors, which is consistent with our *in vivo* observations. Furthermore, the RD-enriched cluster 3 was dominated by TREM2 macrophages, an immunosuppressive tumor-recruited macrophage population that has been shown to suppress T-cell effector functions and express genes related to lipid metabolism and endo-lysosomal pathway.^25–27^ To further characterize the signaling pathways associated with the MoMac clusters, we performed Gene Ontology (GO) biological process enrichment analysis on upregulated differentially expressed genes (DEG) of each cluster (**Figure 4F**). The MoMacs in the TN-enriched cluster 0 expressed signatures of antigen presentation and T cell activation, suggesting pro-inflammatory pathway activation. Cluster 1 was primarily defined by increased cytoplasmic translation and ribosome biogenesis (including rRNA metabolic processing and subunit assembly), a process that in macrophages has been linked to protumorigenic functions.^28^ Cluster 2 was predominantly composed of IL1B monocytes, enriched with proinflammatory processes and chemotaxis, thus resembling monocyte-derived IL1B^+^ TAMs that regulate pathogenic tumor-promoting inflammation within the TME^29^ and consistent with an actively recruited monocyte population within the RD niche. The MoMacs in cluster 3 expressed signatures of lipid metabolism, which is consistent with the lipid-associated TREM2 macrophage phenotype^30–32^ (**Figure 4E, F**). Cluster-specific MoMac DEGs remained predominantly the same, when a confirmatory analysis was conducted without the one PD sample and balanced cell numbers across the samples (**Supplementary Figure S5B**). Together, these results suggest that the MoMacs in the TN tumors are mostly pro-inflammatory, whereas the RD tumors are characterized by the presence of recruited monocytes and immunosuppressive TREM2 macrophages. Indeed, when we directly compared the TN and RD MoMacs in a Hallmark gene set enrichment analysis, we could see a significant suppression of the interferon-alpha signature in the RD MoMacs (**Figure 4G, H**), indicating a suppression of inflammatory signaling in RD MoMacs. We could also see a significant upregulation of the bile acid metabolism -pathway in the RD MoMacs, again consistent with the TREM2 phenotype. While our curated sample set only contained one PD sample, it is noteworthy that all pathways differentially enriched in the RD MoMacs were reverted in the PD MoMacs to resemble the MoMacs in the TN samples (**Figure 4G, H**), suggesting that also in human patients, the changes in the MoMac phenotypes observed in residual tumors are highly specific to the residual state and reversible upon loss of response to EGFR TKI.

Together, these results suggest that in EGFR-mutant lung cancer patients, EGFR TKI treatment leads to widespread changes in monocyte and macrophage composition, driving a treatment-induced switch to an immunosuppressive myeloid tumor microenvironment.

### DTP secretome drives an immunosuppressive TREM2 macrophage phenotype

Building on the observation that osimertinib treatment induces a switch from pro-inflammatory to immunosuppressive TREM2^+^ macrophages in EGFR-mutant lung cancer patients, we examined whether the DTP secretome is sufficient to promote this phenotype switch *in vitro*. For this, we re-analyzed our bulk RNA-seq data from macrophages exposed to DTP-conditioned medium, or conditioned medium from untreated, proliferating cells (**Figure 2F-H**). Performing a GSEA analysis using the MoMacVERSE gene signatures, we were indeed able to see a strong enrichment of the TREM2^+^ macrophage signature, as well as an independently published TREM2 signature^30^, in macrophages exposed to DTP-conditioned medium (**Figure 5A, B**), indicating that secreted factors from DTP cells alone can promote the immunosuppressive TREM2^+^ phenotype in macrophages. Along with the TREM2 signatures, we also observed a significant enrichment of the signature for HES1 macrophages, which have also been associated with immunosuppression^25,33,34^. In contrast, the more proinflammatory ISG^+^ monocyte and IL4l1^+^ macrophage signatures were significantly negatively enriched with exposure to DTP-conditioned medium (**Figure 5A**). Furthermore, GSEA analysis using the Hallmark pathways revealed strong negative enrichment for interferon-alpha, interferon-gamma, and inflammatory response pathways in macrophages exposed to DTP-conditioned medium (**Figure 5C**), demonstrating that the DTP cells drive a highly suppressed proinflammatory program in macrophages, recapitulating our findings from the patient samples (**Figure 4G, H**).

**Figure 5.**
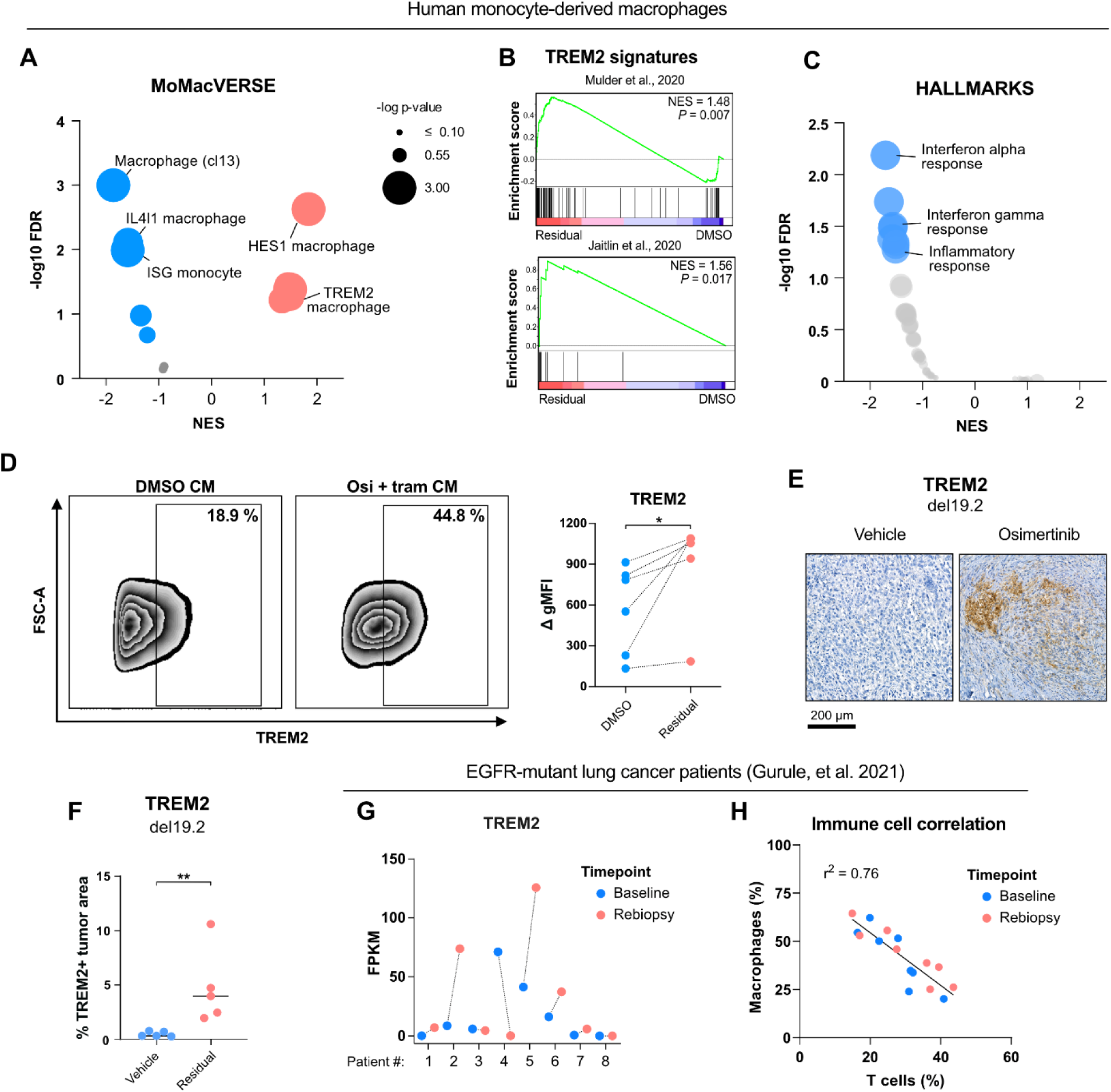
DTP secretome drives an immunosuppressive TREM2 macrophage phenotype. A) GSEA analysis of MoMacVERSE^25^ monocyte/macrophage signatures using RNA-sequencing data from Figure 2F-H. B) GSEA enrichment plots of two independent TREM2^+^ macrophage signatures^25,30^. C) GSEA of Hallmark genes. D) Flow cytometry analysis of TREM2 surface expression in monocyte-derived macrophages exposed to HCC827 CM. E) Immunohistochemical staining of TREM2 from syngeneic Del19.2 model in vivo experiment depicted in Figure 1A-C. F) Quantification of TREM2^+^ area from samples in (E) using QuPath. G) TREM2 gene expression in EGFR TKI-treated patients. RNA-sequencing data from Gurule et al.^35^. H) Correlation of macrophage and T cell abundance from CIBERSORTx analysis of RNA sequencing data in (G). CM = conditioned medium. Paired (D) or unpaired (F) t-tests were used to assess statistical significance. **, P-value <0.01; *, P-value < 0.05.

We could also validate these transcriptional findings at the protein level by flow cytometry, demonstrating that TREM2 surface expression was significantly induced in macrophages exposed to DTP-conditioned medium when compared to conditioned medium from control cells (**Figure 5D**). Furthermore, immunohistochemical analysis of residual tumor samples from the Del19.2 mouse model demonstrated an accumulation of TREM2⁺ macrophages in osimertinib-treated residual tumors (**Figure 5E–F**).

To further evaluate TREM2 expression in EGFR-mutant lung cancer patient samples, we analyzed a published bulk RNA-sequencing dataset from eight patients treated with EGFR TKIs^35^, with matched tumor samples collected before and during treatment. In this cohort, TREM2 expression, almost exclusively restricted to macrophages, increased during therapy in five out of eight patients, suggesting that TREM2 upregulation is a recurrent feature in EGFR TKI-treated tumors (**Figure 5G**). Notably, utilizing CIBERSORT immune cell deconvolution^36^ from the same data set, we observed a strong inverse correlation between macrophage and T cell abundances (**Figure 5H**), suggesting that macrophages may mediate immune evasion in EGFR-mutant lung cancer patients, which aligns with our observations.

Together, these data demonstrate that residual cancer cells can directly promote an immunosuppressive TREM2 macrophage phenotype and suppress pro-inflammatory signaling, shaping an immunosuppressive myeloid microenvironment *via* secreted factors.

### Treatment-induced YAP/TEAD activity in DTP cells drives immunosuppressive reprogramming of macrophages

Activation of Yes-associated protein 1 (YAP) drives resistance to targeted therapy in a wide array of oncogene-addicted cancer types^7,37–40^, including EGFR-mutant lung cancer. Treatment-induced YAP activation has been shown to suppress apoptosis and facilitate a reversible state of dormancy in response to EGFR TKIs, and blocking YAP activity with TEAD inhibitors improves the efficacy of EGFR-targeted therapy *in vitro* and *in vivo* xenograft models of EGFR-mutant lung cancer^7,9,41^. Intriguingly, YAP also has extensive non-cell-autonomous functions^10^, and has been shown to suppress the tumor immune response by driving the expression of soluble factors that recruit macrophages to the tumor microenvironment^11–15^. Therefore, we hypothesized that treatment-induced YAP activity could also contribute to the DTP-driven macrophage phenotype switch through YAP-driven expression of secreted factors.

To assess the contribution of YAP activity to the DTP secretome, we treated HCC827 and HCC4006 cells for 10 days with osimertinib/trametinib to induce the stable residual state, followed by a 48-hour treatment with TEAD inhibitor MYF-01-37^7^, and performed bulk RNA-sequencing (**Figure 6A**). Using a curated list of genes encoding human secreted proteins^42^, we found that in both cell lines, more than 400 protein-coding secretome genes were significantly upregulated, and nearly 200 downregulated in DTP cells (**Figure 6B**). Of the genes upregulated in DTPs, approximately 25% were regulated by YAP/TEAD in both cell lines, with a significant overlap in YAP/TEAD-driven secretome components (**Figure 6C**). Notably, 56% (25 out of 45) of the shared YAP/TEAD -driven secretome genes have been previously linked to monocyte/macrophage activation, recruitment, and/or polarization (**Figure 6F, Supplementary Table S2**), indicating that the YAP/TEAD -driven secretome is rich in factors regulating monocyte/macrophage biology. Utilizing our previously published PC-9 YAP and TEAD4 ChIP-seq data^7^, we could confirm the direct binding of YAP and TEAD to the genomic loci of 38 out of 45 (84 %) shared YAP/TEAD-driven secretome genes upon 48-hour osimertinib/trametinib treatment (**Figure 6E, Supplementary Figure S6A**), suggesting that YAP is directly regulating the expression of these genes through TEAD. Furthermore, by performing mass spectrometry analysis from the conditioned medium of residual cancer cells with or without MYF-01-37 treatment, as well as from DMSO-treated cells, we could confirm in protein level the increased secretion of 17 out of 45 shared YAP/TEAD -driven secretome genes (**Supplementary Figure S6B, C**).

**Figure 6.**
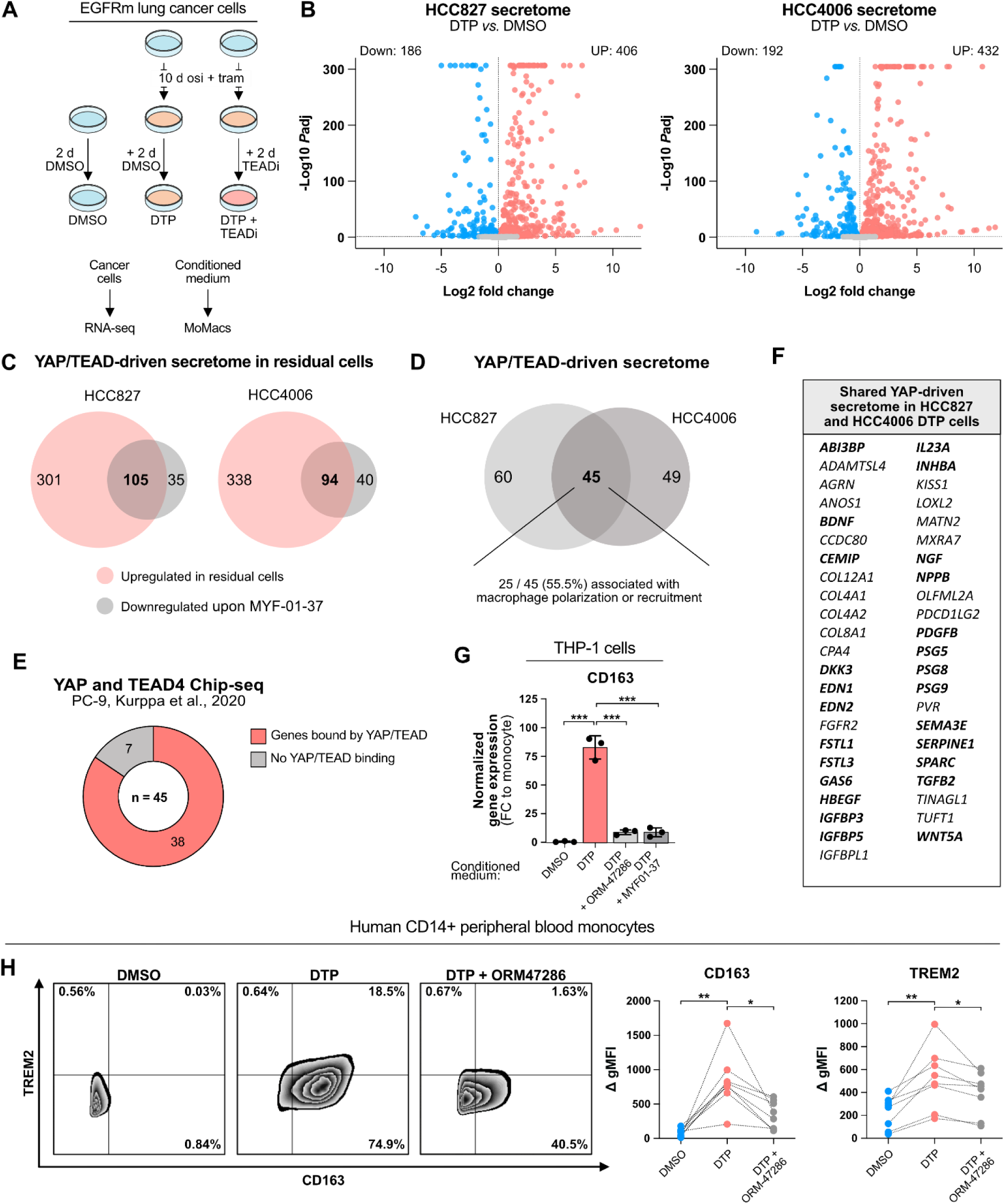
Treatment-induced YAP/TEAD activity in DTP cells drives immunosuppressive reprogramming of macrophages. A) Experimental set-up for conditioned media collection and YAP/TEAD inhibition in DTP cells. B) RNA-sequencing analysis of the expression of genes encoding for secreted proteins in HCC827 and HCC4006 cell lines treated as in (A). C) YAP-driven secretome in HCC827 and HCC4006 cells, determined by genes upregulated in DTPs, and downregulated upon TEAD inhibition. D) Shared YAP-driven secretome in HCC827 and HCC4006. E) ChIP-seq analysis of YAP binding of shared secretome genes. F) List of shared YAP-driven secretome from (D), with genes previously associated with macrophage activation, migration, or polarization highlighted in bold. G) CD163 expression of THP-1 cells treated with HCC827 conditioned media collected as in (A) using 500 nM ORM-47286 or 10 mM MYF-01-37 for TEAD inhibition. H) Flow cytometry analysis of CD163 and TREM2 cell surface expression in human peripheral blood monocytes exposed to HCC827 conditioned media. Data in (G) presented as mean ± SD. Unpaired (G) and paired (H) one-way ANOVA was used to assess statistical significance. **, P-value <0.01; *, P-value < 0.05.

To evaluate the functional impact of YAP/TEAD -driven secretome on DTP-driven monocyte reprogramming, we collected conditioned medium from HCC827 cells treated identically as for the RNA-seq experiment and depicted in **Figure 6A**, but including two different, structurally divergent TEAD inhibitors; MYF-01-37 and a pan-TEAD inhibitor ORM-47286^43^. Stimulating THP-1 monocytes with these conditioned media, we observed a significant dampening of DTP-driven *CD163* induction in THP-1 cells when the DTP cells were treated with either of the TEAD inhibitors (**Figure 6G**), indicating that YAP/TEAD activity in DTP cells is driving the immunosuppressive reprogramming of THP-1 cells. In contrast, applying the TEAD inhibitors directly to THP-1 monocytes resulted in only a modest reduction in M2 marker induction (**Supplementary Figure SDC**), confirming that the monocyte reprogramming arises from YAP/TEAD activity in the cancer cells rather than from direct effects on monocytes. Importantly, we could also replicate these observations in primary human CD14^+^ monocytes using flow cytometry, where the treatment of DTP cells with ORM-47286 blunted the DTP-driven induction of both CD163 and TREM2 surface expression (**Figure 6H**), suggesting that the residual state -driven immunosuppressive TREM2-phenotype we observed *in vitro*, *in vivo*, and in human patients, is driven by YAP/TEAD activity in the residual cancer cells.

Together, these results demonstrate that YAP/TEAD activity in the DTP cells drives the expression and secretion of soluble factors that promote an immunosuppressive TREM2 macrophage phenotype in human MoMacs.

### TEAD inhibition decreases immunosuppressive macrophage infiltration and enhances treatment efficacy in syngeneic mouse models of EGFR-mutant lung cancer

Our observation that YAP/TEAD inhibition in DTP cells reduces their ability to drive immunosuppressive priming and TREM2 expression in THP-1 cells and human primary monocytes suggests that targeting YAP/TEAD activity in combination with osimertinib may have a dual effect; preventing both cell-intrinsic and microenvironment-mediated mechanisms of resistance. To assess whether co-inhibition of YAP/TEAD activity could also remodel the residual tumor microenvironment *in vivo* and enhance the efficacy of EGFR TKI therapy in fully immunocompetent mice, we tested the combination of osimertinib and ORM-47286 in the syngeneic del19.2 and L860R mouse models. While ORM-47286 alone had minimal impact on tumor growth, the combination of ORM-47286 with osimertinib led to a significant decrease in residual cancer cell burden and significantly prolonged the time to relapse following treatment cessation in both models (**Figure 7A–C**). Notably, in the del19.2 model, the osimertinib + ORM-47286 combination led to almost complete eradication of the tumors, with only a single mouse relapsing after treatment cessation in the combination arm, compared to a more widespread relapse in the osimertinib-only cohort (7 out of 20 tumors).

**Figure 7.**
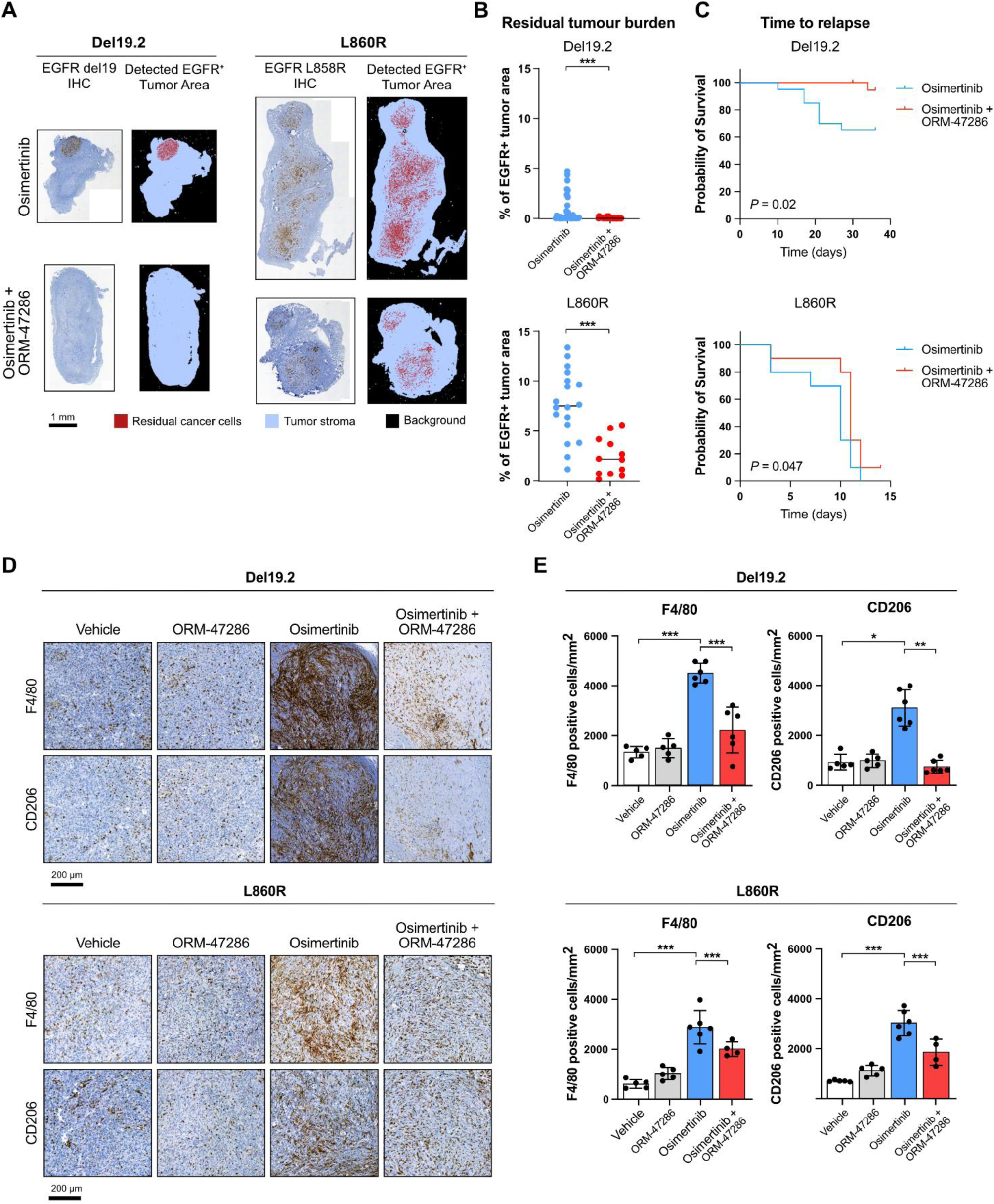
TEAD inhibition decreases immunosuppressive macrophage infiltration and enhances treatment efficacy in syngeneic mouse models of EGFR-mutant lung cancer. A) Determining residual cancer cell burden in Del19.2 and L860R residual tumors following treatment with osimertinib or the combination of osimertinib and ORM-47286. Sections stained with EGFR Del19 or EGFR L858R -specific antibody were analyzed using artificial neural network-based pixel classification in QuPath. B) Quantification of residual cancer cell burden using analysis in (A), determined by either EGFR Del19 or L858R positive tumor area (n = 5-7 slides/tumor, 6 mice/cohort). C) Relapse-free survival of treated mice, determined as time to tumor regrowth after treatment cessation to the original size at day 0. E) Immunohistochemical staining of tumor-infiltrating macrophages in samples from (A). F4/80, macrophages; CD206, M2 macrophage marker. C) Quantification of F4/80 or CD206 -positive cells per mm^2^ in tumor sections depicted in (E) using QuPath. Data in (F) presented as mean ± SD. Mann-Whitney test (C), Log-rank (Mantel-Cox) test (D), and one-way ANOVA (F) was used to assess statistical significance. ***, P-value <0.001; **, P-value <0.01; *, P-value < 0.05.

We further investigated the impact of YAP/TEAD inhibition on osimertinib-induced immunosuppressive macrophage accumulation using immunohistochemistry from residual tumors after three weeks of treatment. We observed that the combination of osimertinib and ORM-47286 significantly reduced the accumulation of F4/80^+^ macrophages and, critically, decreased the number of CD206^+^ immunosuppressive macrophages in residual tumors across both models, when compared to osimertinib monotherapy (**Figure 7D, E**). These data suggest that YAP/TEAD activity in osimertinib-treated tumors promotes the accumulation of immunosuppressive macrophages *in vivo*, and this can be reversed by concomitant TEAD inhibition, in agreement with our *in vitro* results. Taken together, these data suggest that in fully immunocompetent models of EGFR-mutant lung cancer, YAP/TEAD inhibition not only enhances the efficacy of EGFR TKI therapy but also alters the myeloid landscape of residual tumors to a less immunosuppressive state, thus potentially suppressing immune evasion and relapse.

## Discussion

EGFR-targeted therapy leads to favorable initial responses in EGFR-mutant NSCLC, yet acquired resistance remains inevitable. Following initial response, drug-tolerant cells can persist in a reversible, quiescent state in residual tumors before eventually giving rise to a drug-resistant tumor^44^. Treatment-naïve EGFR-mutant tumors are typically immunologically “cold” and do not benefit from immune checkpoint inhibition.^45^ Paired-sample studies comparing treatment-naïve EGFR-mutant NSCLC to post-EGFR TKI resistance biopsies often report reduced infiltration of CD8^+^ T cells and increased PD-L1 expression^46–48^, consistent with an immunosuppressive shift. Thus, targeted therapy-induced changes in immune composition during treatment could potentially shape both the depth of response and the trajectory to resistance.

It is widely acknowledged that oncogene-targeted therapy induces broad transcriptional and epigenetic rearrangements in surviving cancer cells, leading to highly plastic DTP cell states. Intriguingly, these states are associated with activation of cancer cell-intrinsic inflammatory response, reversible senescence program, and increased expression of genes encoding for secreted proteins^3–6^, raising the question whether the phenotypically altered DTP cells could take part in shaping the tumor microenvironment and, particularly, modulate the tumor immune response in on-treatment tumors. Here, we set out to investigate the contribution of on-treatment DTP cells to the tumor immune response and show that in the context of EGFR-mutant lung cancer, DTP cells actively remodel the tumor immune microenvironment during treatment. The quiescent, on-treatment cancer cells exploit a YAP/TEAD-driven secretome to recruit and reprogram monocytes into immunosuppressive M2-like, TREM2^+^ macrophages with suppressed proinflammatory interferon signaling and the ability to blunt T-cell effector functions. We further demonstrate that interrupting this axis with TEAD inhibition reduces the infiltration of immunosuppressive macrophages and extends therapeutic responses in immunocompetent mouse models.

We observed that the ability of residual DTP cells to reprogram macrophages to the immunosuppressive M2-like/TREM2^+^ phenotype was highly dependent on a stable quiescent state, both *in vitro* and *in vivo*. EGFR-mutant lung cancer cells treated with osimertinib *in vitro*, which re-initiate cell cycle within a few days on treatment^7,18,19^ (**Figure 2B**), were inferior in promoting the M2-like phenotype in THP-1 cells compared to cells treated with the combination of osimertinib and MEK inhibitor trametinib. This combination prevents the reactivation of the MAPK pathway and leads to a stable quiescent state^7,18,19^ (**Figure 2C**). Analogously, treatment of PC-9 xenografts *in vivo* with a suboptimal osimertinib dose led to less prominent accumulation of immunosuppressive macrophages than optimal dosing with a sustained response (**Figure 1D–F)**. Consistently, osimertinib/trametinib-treated DTP cells exposed to a drug holiday lost the ability to reprogram THP-1 cells (**Figure 2D)**. In the syngeneic mouse models, cessation of osimertinib treatment and subsequent tumor regrowth led to reversal of the macrophage accumulation observed in residual, on-treatment tumors, to levels resembling those in vehicle-treated mice (**Figure 1A-C**). Finally, in EGFR-mutant lung cancer specimens, macrophages in the progressive disease sample were phenotypically similar to those in samples from treatment-naïve patients, and did not exhibit the M2-like/TREM2^+^ phenotype observed in macrophages of the residual, on-treatment tumors (**Figure 4**). Consistently, Fang *et al* reported an increase in overall immune scores after EGFR TKI, but found no significant changes between treatment-naïve and TKI-resistant samples.^49^ Together, these observations suggest that quiescent DTP cells have a particular capability to remodel the myeloid microenvironment in residual tumors, and that this capability is lost upon relapse. This, in turn, raises the possibility that residual tumors may harbor novel immunotherapy opportunities which may be difficult, if not impossible, to predict from treatment-naïve tumors, and may not persist in relapsed tumors.

Our study focused on macrophages due to the striking accumulation of M2-like/TREM2^+^ macrophages in residual tumors following osimertinib treatment in both syngeneic mouse models and in patient samples. Macrophages are increasingly recognized as an attractive immunotherapy target across various cancer types due to their capacity to suppress the tumor immune response and support tumor growth.^50,51^ Consistently, our results demonstrate that the DTP-reprogrammed monocytes strongly suppress T-cell activation *in vitro* (**Figure 3**), and that macrophage abundance inversely correlates with T-cell infiltration in EGFR-mutant lung cancer patient samples (**Figure 5H**), in line with findings from an independent cohort of EGFR-mutant lung cancer patients.^52^ These results support the view that treatment-induced, DTP-driven infiltration and reprogramming of M2-like/TREM2^+^ macrophages is immunosuppressive. As M2-like macrophages have been previously associated with poor response to EGFR TKIs^53^, and T-cell response has been shown to be crucial for sustained EGFR TKI response in the Del19.2 syngeneic mouse model^16^, it is plausible that the treatment-induced accumulation of immunosuppressive M2-like/TREM2^+^ macrophages helps sustain cancer cell survival in residual tumors by facilitating immune evasion.

This raises the question of whether direct interference with macrophage accumulation and the subsequent formation of an immunosuppressive niche could enhance the efficacy of targeted therapy. Indeed, several studies have explored strategies to deplete or reprogram macrophages in a cancer context^50^. We therefore explored the possibility of enhancing the efficacy of osimertinib by depleting macrophages using a CSF1R inhibitor pexidartinib. Unexpectedly, pexidartinib preferentially depleted the MHCII^high^ (M1-like) macrophages while having little effect in depleting MCHII^low^ (M2-like) macrophages (**Supplementary Figure S3**), leading to decreased T-cell infiltration and no benefit over osimertinib alone. Our results are contrary to Okawa *et al*, who recently showed in a syngeneic EGFRdel19 model that the addition of pexidartinib provides a modest additional benefit over osimertinib treatment alone by reducing CD206^+^ macrophage accumulation^54^. While we cannot exclude the possibility that experimental design or different EGFR driver mutation contexts could result in opposing outcomes in pexidartinib combination therapy, we find this unlikely. Rather, our data underscore the context dependence and highlight the risk of CSF1R-targeted macrophage depletion backfiring in different preclinical models, and potentially in patients, with indiscriminate macrophage depletion particularly when MHCII^high^ (M1-like) populations are preferentially removed, skewing the M1/M2 balance and lowering CD8^+^ T cell infiltration.

As these findings suggest that macrophage-depleting interventions may be insufficient to reverse the immunosuppressive state established during EGFR-targeted therapy, we turned to the cancer cell-intrinsic mechanisms that drive the formation of the immunosuppressive macrophage niche. Given the established link between YAP activation, DTP cell survival, and EGFR TKI resistance^7,41^, as well as its increasingly apparent role in regulating tumor immune cell infiltration and antitumor immunity^12–15,55–58^, we hypothesized that YAP/TEAD-dependent transcriptional programs may orchestrate the immunomodulatory secretome that shapes the residual tumor microenvironment. Indeed, we observed that beyond enabling DTP cell survival following targeted therapy, sustained YAP activation in DTP cells drives a broad secretome enriched for macrophage-modulating factors (**Figure 6B–F**). Functionally, TEAD inhibition in DTP cells (but not in monocytes) significantly attenuated the expression of immunosuppressive markers in THP-1 cells and human primary monocytes (**Figure 6G, H, Supplementary Figure S6D**). *In vivo*, TEAD inhibition reduced F4/80^+^/CD206^+^ macrophage accumulation in the residual tumor microenvironment (**Figure 7E, F**), and deepened osimertinib response and delayed relapse (**Figure 7A–D**). Collectively, our findings position YAP/TEAD as an important driver of an immunosuppressive TME in on-treatment tumors. Our findings suggest that through EGFR TKI-induced quiescence with a YAP-driven secretory profile, DTP cells can effectively remodel the surrounding microenvironment to favor immune evasion. Our results further suggest that by targeting TEAD, tumor cell secretory outputs can be rewired to limit the formation of an immunosuppressive niche while simultaneously targeting tumor cell-intrinsic survival signaling, thus potentially counteracting both cancer cell-intrinsic and microenvironmental mechanisms of resistance. As TEAD inhibitors hold great promise in the combination setting in various oncogene-addicted cancer contexts^8,9^, and are migrating towards clinical studies in this setting, our hypothesis should be evaluated in EGFR-mutant lung cancer patients undergoing TEAD inhibitor combination therapy in the future.

Together, our results are consistent with a paradigm in which treatment-induced plasticity of cancer cells gives rise to drug tolerant phenotypes that can directly drive immunosuppressive changes in the tumor microenvironment, with these changes being specific to sustained therapeutic response and reversible upon relapse. This may have several consequences: 1) strategies to enhance the immune response in EGFR-mutant lung cancer patients treated with EGFR-targeted therapies should be derived through understanding the changes in the TME that occur *during* treatment, rather than by analyzing treatment-naïve or relapsed tumors, 2) treatment-induced changes may provide novel, residual state -specific opportunities for immunotherapy that may not exist in the treatment-naïve or relapsed tumors, and 3) as the on-treatment, residual cancer cells are active drivers of TME modulation in on-treatment tumors, targeting the key cell-intrinsic nodes promoting this modulation, such as YAP/TEAD, present attractive therapeutic targets for preventing treatment-induced adaptive immunosuppression, potentially prolonging the efficacy of oncogene-targeted therapies.

## Materials and methods

### Cell culture

EGFR-mutant lung cancer cell lines HCC827, HCC4006, and PC-9 cells were cultured in complete RPMI-1940 media (EuroClone, cat# ECB9006L) supplemented with 10 % FBS (Gibco, cat# A5256701), 2 mM L-glutamine (EuroClone, cat# ECB3004D), and 1 U penicillin/streptomycin (EuroClone, cat# ECB3001D). THP-1 monocytic cell line was cultured in full IMDM media (Gibco) with 10 % FBS and penicillin/streptomycin. Syngeneic mouse EGFR-mutant lung cancer cell lines Del19.2 and L860R were generously gifted by Lynn E Heasley and Raphael Nemenoff.^16^ Mouse cell lines were cultured in RPMI-1940 supplemented with 5 % FBS, L-glutamine, and penicillin/streptomycin. The Del19.2 and L860R cells harbor a mutantEgfr-2A-mCerulean3 transgene. To avoid potential immunogenicity of the mCerluean3, the mCerulean3 was knocked out from both cell lines using CRISPR/CAS9 as previously described^7^, using a guide sequence TTCAAGTCCGCCATGCCCGA, and the EN-138 nucleofection program in the Lonza 4D Nucleofector. The mCerulean3 negative populations were sorted 5 days after nucleofection by FACS, expanded, and used in the experiments.

### Animal experiments

All animal experiments were performed per the Finnish Act on Animal Experimentation (62/2006) and were approved by the Committee for Animal Experimentation (license number ESAVI/7740/2023, ESAVI/39706/2021). For PC-9 xenograft experiments, PC-9 cells were detached, washed with PBS, and resuspended in 1:1 serum-free RPMI-1940 and Matrigel (Corning, cat# 354234). 7 to 8-week-old female Athymic Nude-Foxn1nu (Envigo) mice were subcutaneously inoculated with 5 x 10^6^ cells / flank. Once tumors reached average size of 300 mm^3^, the mice were randomly assigned to receive daily doses of either vehicle (0.5% methylcellulose/0.5% Tween80) or 2.5 or 10 mg/kg osimertinib by oral gavage (6 to 10 mice / group). Mice were monitored daily, and tumors were measured three times a week using a caliper. At the specified timepoints, the mice were sacrificed, and the tumors were fixed in 4 % PFA for 48 hours, transferred to 70% ethanol and embedded in paraffin. For experiments with syngeneic EGFR-mutant lung cancer models, Del19.2 and L860R cells were detached, washed once with PBS, and resuspended in 1:1 phenol red-free DMEM (Gibco, cat# 11520556) and Matrigel. 6 to 8-week-old male C57BL/6J mice were subcutaneously inoculated with 5 x 10^5^ cells/flank. Mice were monitored daily, and tumors were measured three times a week using a caliper. Once tumors reached an average size of 250-400 mm^3^, mice were randomized into treatment groups (5 to 10 mice per group). Osimertinib was dosed in mouse diet containing 35mg/kg osimertinib (ssniff Spezialdiäten GmbH). For combination treatments, mice were dosed with osimertinib in the diet, and either 200 µg i.p. injections of mAbMods™ anti-Mouse TREM2 (CliniSciences, cat# T721-5) or its isotype control IgG (CliniSciences, cat# I-1241-5) every five days, daily 400 µg pexidartinib (MedChemExpress, cat# HY-16749) in 10% DMSO, 0.5% methylcellulose/0.5% Tween80 (Thermo Scientific Chemicals) by oral gavage or daily 10 mg/kg ORM-47286 (Orion Pharma) in 0.5% methylcellulose/0.5% Tween80 by oral gavage (Thermo Scientific Chemicals). Tumor samples were harvested and processed as the PC-9 tumors.

### Immunohistochemistry and quantitation

Immunohistochemical staining was performed on paraformalin-fixed, paraffin-embedded tumors. The sections were deparaffinized, rehydrated, and underwent antigen retrieval using either citrate buffer (pH 6) or Tris-EDTA buffer (pH 9). Endogenous peroxidase activity was inhibited with hydrogen peroxide, and nonspecific staining was blocked using BrightDiluent normal antibody diluent (BD09-125, Immunologic, Arnhem, Netherlands). All antibodies were purchased from Cell Signaling Technologies (Danvers, MA, USA) unless otherwise specified. Primary antibodies used in staining were EGFR E746-A750del Specific (clone D6B6, 1:400, cat#2085), EGFR L858R Mutant Specific (clone 43B2, 1:100, cat#3197), F4/80 (clone D2S9R, 1:500, cat#70076), Arginase-1 (clone D4E3M™, 1:400, cat#93668), CD206 (clone E6T5J, 1:400, cat#24595), TREM2 (clone EPR26210, 1:200, cat#305103, abcam, Cambridge, UK), and CD8α (clone D4W2Z, 1:400, cat#98941). Primary antibody staining was carried out for 60 minutes at room temperature. Sections were stained with Lab Vision™ autostainer (ThermoFisher Scientific). Primary antibodies were visualised with horseradish peroxidase (HRP)–conjugated secondary antibodies (BrightVision goat anti-rabbit IgG HRP detection system, cat#DPVR110HRP, Immunologic). Slides were counterstained with Meyer’s hematoxylin for one minute at room temperature. Stained sections were digitalized using a P1000 slide scanner (3DHISTECH, Budapest, Hungary) and quantified using QuPath 0.4.4 software^59^ as positive cells/mm^2^. Residual cancer cell burden was quantified from EGFR IHC-stained 3–7 whole-slide scans. A pixel classifier based on an artificial neural network (ANN_MLP) was trained on 228 annotated tumor regions from 18 mouse samples included in the experiments shown in Figure 1 (Del19 and L860R models). The classifier used Gaussian, Laplacian, and weighted_std_dev features derived from the hematoxylin, HDAB, and residual image channels. Regions of interest (ROIs) were manually annotated to exclude non-tumor tissue, and classifier output was manually reviewed for accuracy. The trained model was applied to subsequent in vivo experiments, and EGFR-positive tumor burden was reported as a percentage of the total annotated tumor area.

### Cell cycle analysis

HCC827 and PC-9 cells lentivirally transduced with pLEX307-FUCCI cell cycle reporter were seeded at 4000 cells/well on a 96-well plate. 24 hours after seeding, the media was replaced, and treatment with 100 nM osimertinib ± 30 nM trametinib was started. Cells were imaged every two hours for up to 24 days using the Incucyte S3 live cell imaging microscope. Media was changed every 3-5 days. For quantitation, 3 images were taken per well to determine cell confluency and cell cycle state. G1 and G1/S/M states were determined as red and green cell counts, respectively.

### Flow cytometry from mouse tumors

Tumors were dissociated into single-cell suspensions using the Mouse Tumor Dissociation Kit (Miltenyi, cat# 130-096-730), followed by immune cell enrichment with Mouse CD45 MicroBeads (Miltenyi, cat# 130-052-301). Tumor-associated macrophages and tumor-infiltrating lymphocytes were stained using two separate antibody panels. Fixable Viability Dye eFluor™ 450 or 780 (Invitrogen, eBioscience™, 1:1000) was used for viability assessment, and Mouse BD Fc Block™ (Purified Rat Anti-Mouse CD16/CD32, clone 2.4G2, BD Pharmingen™, cat# 553141, 1:50) to prevent unspecific antibody binding.

Surface staining for markers was carried out in PBS + 2 % FBS for the following antibodies (1:100 dilution)CD45-AF700 (clone 30-F11, Invitrogen, eBioscience™), CD11b-APC-Cy7 (clone M1/79, BD Pharmingen™), Ly-6G-AF700 (clone 1A8, BD Pharmingen™), Ly-6C-PE-Cy7 (clone HK1.4, Invitrogen, eBioscience™), MHC II-PerCP-Cy5.5 (clone M5/144, BD Pharmingen™), CD206/MMR-PE (clone C068C2, Biolegend), CD3-PerCP-Cy5.5 (clone 17A2, BD Pharmingen™), CD4-BV510 (clone GK1.5, Biolegend), and CD8a-BV650 (clone 54-6.7, BD Horizon™). For compensation controls, individual fluorophores were recorded using single-stained UltraComp eBeads™ (Thermo Fisher Scientific, cat# 1-2222-42 at a final concentration of 1:100). Data acquisition was performed on a BD LSRFortessa™ cytometer and analyzed using FlowJo™ software (BD Biosciences) (**Supplementary Figure 4E**).

### Collection of cancer cell-conditioned media

To generate drug-tolerant residual cells, human 6-10×10^5^ EGFR-mutant NSCLC cell lines were seeded on 10 cm cell culture plates and treated with 100 nM osimertinib (Cayman Chemical, cat# 16237 or MedChemExpress, cat# HY-15772) with or without 30 nM trametinib (MedChemExpress, cat# HY-10999) for ten days in full RPMI medium, with medium and drugs refreshed after 5 days of treatment. 1-5×10^5^ cells were plated as control 24 hours before collecting cancer-cell conditioned media. To condition media on both proliferating and quiescent residual cells, cells were washed once with PBS, followed by incubation in full IMDM media with 100 nM osimertinib + 30 nM trametinib or DMSO (Fisher Scientific, cat# BP231-100). After 48 hours of incubation, cancer cells were trypsinized and counted while conditioned media (CM) was collected and filtered through 0.45 µg syringe filter and stored in -80°C if not used fresh.

### RNA-sequencing of cancer cells

2.5 x 10^6^ HCC827 cells and 1.5 x 10^6^ HCC4006 cells were plated on 15 cm cell culture plates for DTP generation, and treated for 10 days with 100 nM osimertinib + 30 nM trametinib, followed by addition of 10 µM MYF-01-37 or DMSO for further 48 hours. Control cells (8 x 10^5^ cells/plate) were treated with DMSO for 48 hours. Cells were collected, counted, and lysed for RNA isolation using NucleoSpin® RNA purification kit (Macherey-Nagel cat#740955) according to manufacturer’s instructions. 100 ng of RNA was used for library preparation. RNA quality was confirmed at the Finnish Functional Genomics Centre (FFGC), and libraries were prepared using the Illumina Stranded mRNA Preparation kit with poly(A) selection. Libraries were sequenced on an Illumina NovaSeq 6000. Differences in gene expression of distinct treatments and cell types were assessed by DESeq2^60^ using a multifactor design for paired analysis.

### Multiplex secretome analysis from cancer-cell conditioned media

Cytokine levels were measured using Bio-Plex Pro Human Cytokine 27-plex assay (Bio-Rad, cat. M500KCAF0Y) and Bio-Plex 200 System (Bio-Rad) according to the manufacturer’s instructions. Multiplex readouts were normalized to cell count. Results were presented as fold changes between DMSO and osimertinib + trametinib-treated samples.

### Proteomics analysis from cancer-cell conditioned media

Secretome enrichment from cell culture media was performed with ProteoMiner kit (Bio-Rad) according to manufacturer’s instructions. The enriched proteins were reduced with 10 mM D,L-dithiothreitol and alkylated with 40 mM iodoacetamide, and digested overnight with sequencing grade modified trypsin (Promega) in 1M urea, 50mM Tris. After digestion peptide samples were desalted with a Sep-Pak tC18 96-well plate (Waters) and evaporated to dryness.

The LC-ESI-MS/MS analyses were performed on a nanoflow HPLC system (Easy-nLC1200, Thermo Fisher Scientific) coupled to the Orbitrap Fusion Lumos Tribrid mass spectrometer (Thermo Fisher Scientific, Bremen, Germany) equipped with a nano-electrospray ionization source and FAIMS interface. FAIMS Compensation voltages of -50 V and -70 V were used. Peptides were first loaded on a trapping column and subsequently separated inline on a 15 cm C18 column (75 μm x 15 cm, ReproSil-Pur 3 μm 120 Å C18-AQ, Dr. Maisch HPLC GmbH, Ammerbuch-Entringen, Germany). The mobile phase consisted of water with 0.1% formic acid (solvent A) or acetonitrile/water (80:20 (v/v)) with 0.1% formic acid (solvent B). A 120 min step gradient (from 5 to 21% of solvent B in 60 mins, to 36% of solvent B in 48 mins, to 100% of solvent B in 6 min, followed by a 6 min wash stage with 100% of solvent B) was used to eluate peptides.

Samples were analysed by a data independent acquisition (DIA) LC-MS/MS method. MS data was acquired automatically by using Thermo Xcalibur 4.1 software (Thermo Fisher Scientific). In a DIA method, a duty cycle contained one 120000 resolution full scan (400-1000 m/z) and 40 DIA MS/MS scans at 30000 resolution covering the mass range 400-1000 with variable width isolation windows. Data analysis consisted of protein identifications and label free quantifications of protein abundances. Data was analyzed by Spectronaut software (Biognosys; version 17.3). DirectDIA approach was used to identify proteins and label-free quantifications were performed with MaxLFQ. Main data analysis parameters in Spectronaut: Enzyme: Trypsin/P; Missed cleavages: 2; Fixed modifications: Carbamidomethyl*; Variable modifications: Acetyl (protein N-term) and oxidation (M); Protein database: Swiss-Prot 2023_01 Homo Sapiens, Trembl 2023_01 Bos Taurus, Universal Protein Contaminant database^61^ Precursor FDR Cutoff: 0.01; Protein FDR Cutoff: 0.01; Quantification MS level: MS2. Quantification type: Area under the curve within integration boundaries for each targeted ion; Normalization: Local normalization (Based on RT dependent local regression model described by Callister et al., 2006.^62^

### Isolation of monocytes from EDTA blood and generation of macrophages in vitro

Peripheral blood mononuclear cells were isolated from EDTA blood collected from healthy blood donors, using Ficoll-Paque™ PLUS (Cytiva, cat# 17144002) gradient centrifugation. Monocytes were further isolated by positive magnetic selection using MACS LS-columns (Miltenyi, cat# 130-042-401) and human CD14 MicroBeads (Miltenyi, cat#130-050-201) per manufacturer’s instructions. To differentiate isolated monocytes into macrophages, freshly isolated cells were cultured in full IMDM media supplemented with 50 ng/ml recombinant human M-CSF (Biolegend, cat#574804) for six days.

### Polarization of monocytes and macrophages using cancer cell-conditioned media

Human monocytes and macrophages were obtained as described above. THP-1 monocytes or human peripheral blood monocytes and monocyte-derived macrophages were plated either 1-2×10^6^ cells/well on 6-well or 2×10^5^ cells/well on 96-well ultra-low attachment plates (Corning™, Fischer Scientific) in cancer cell-conditioned media collected from cancer cells. Monocytes and macrophages were incubated in CM for 6 days in total. The polarization of monocytes and macrophages was determined using RT-qPCR, flow cytometry, and RNA sequencing.

### Gene expression analysis with RT-qPCR

THP-1 cells were washed once with PBS, detached by incubation in 10 mM EDTA-PBS followed by gentle scraping, centrifugated, lysed for RNA isolation using NucleoSpin RNA isolation kit (Macherey-Nagel, cat# 740955) followed by reverse transcribing 1 µg of RNA/sample to cDNA with SensiFAST cDNA synthesis kit (Bioline, cat# BIO-65054) following manufacturers’ instructions. Gene expression levels were determined with quantitative PCR using QuantStudio 3™ machinery with a hold 2 min 50°C, 10 min 95°C, followed by 40 cycles of 15 sec 95°C and 60 sec 60°C. The following reagents were used to determine gene expression levels; Taqman 2X Universal master mix (cat# 4440047), and 20X gene expression assays GAPDH (Hs02758991_g1), ACTB (Hs99999903_m1), TBP (Hs00427620_m1), MRC1/CD206 (Hs00267207_m1), and CD163 (Hs00174705_m1). Relative gene expression levels were determined using the ΔΔCq data analysis method normalised to at least two separate reference genes/sample.

### Flow cytometry for macrophage polarization

Macrophage polarization for flow cytometry was performed as described previously. Cell viability was assessed with Fixable Viability Dye eFluor™ 450 (Invitrogen, eBioscience™, 1:1000), and Human BD Fc Block™ (clone Fc1, BD Pharmingen™, cat# 564219, 1:50) was used to diminish nonspecific antibody binding. Surface staining for markers was carried out in PBS + 2 % FBS with the following antibodies (1:100 dilution): CD64-PE (clone 10.2, BD Pharmingen™), CD206-AF488 (clone 15-4, BioLegend), CD163-BV711 (clone GHI_61, BD Horizon™), and TREM2-APC (clone 237920, R&D Systems). Corresponding isotype controls APC-Rat IgG2b (clone A95-1, BD Pharmingen™), BV711-Mouse IgG1 (clone X40, BD Horizon™), Alexa Fluor® 488-Mouse IgG1 (clone MOPC-21, BD Pharmingen™) were included to assess nonspecific antibody binding. Gating strategy to assess monocyte/macrophage surface marker expression is shown in **Supplementary Figure 4E**.

### RNA-sequencing of human MoMacs

To analyse the gene expression changes in human monocytes and monocyte-derived macrophages, CD14^+^ peripheral blood monocytes were collected from four volunteer blood donors. 1×10^6^ monocytes and monocyte-derived macrophages were polarised using cancer-cell conditioned media as described previously. After 6 days of incubation, cells were harvested and lysed for RNA isolation using NucleoSpin® RNA purification kit according to manufacturer’s instructions. 300 ng of RNA was used for mRNA sequencing at Novogene (Cambridge, United Kingdom).

### Co-culture of monocytes and T cells

Peripheral blood monocytes were isolated from three healthy blood donors using the Miltenyi Pan Monocyte Isolation Kit (Miltenyi Biotec, cat#130-096-537, human). Isolated monocytes were conditioned for six days in HCC827-conditioned media. One day prior to co-culture, T cells were isolated from matched donors using the Naive Pan T Cell Isolation Kit (Miltenyi Biotec, Cat. 130-097-095, human) and subsequently activated using plate-bound anti-CD3 antibodies. After the monocyte conditioning period, the conditioned media was removed, and T cells were added to the monocytes (1:1 Monocyte to T cell ratio). The co-culture was maintained for an additional 72 hours before flow cytometry analysis. Cell proliferation was measured by pre-staining T cells with CellTrace™ Violet (Thermo Fisher, cat#C34557). For activation analysis, cells were first incubated with BD GolgiStop™ (BD Biosciences, cat#554724) for 6 hours. Cells were then stained for live/dead discrimination in Phosphate-Buffered Saline (PBS) (1:500 dilution for L/D). Surface markers were stained in FACS buffer (5% Fetal Bovine Serum and 2mM EDTA) using anti-CD3 (KIRAVIA Blue 520™; Biolegend, cat#300482), anti-CD4 (Brilliant Violet 605™; Biolegend, Cat. 317438), anti-CD8 (Brilliant Violet 510™; Biolegend, cat#344732), and anti-CD45RO (PE/Cyanine7; Biolegend, Cat. 304230). This was followed by the BD Perm/Wash™ procedure (BD Biosciences, cat#554723) for intracellular staining with anti-Granzyme B (PE; Thermo Fisher, Cat. MHGB04), anti-IFN-γ (Alexa Fluor® 647; Biolegend, cat#502516), and anti-TNF (PerCP/Cyanine5.5; Biolegend, cat#502926). Antibody dilutions were: 1:200 for anti-CD3; 1:100 for all others. Samples were acquired on The NovoCyte Quanteon 4025 and analyzed for mean fluorescence intensity (MFI) and cell proliferation (CellTrace Violet dilution).

### Single-cell RNA sequencing analyses

Single-cell RNA-sequencing data from advanced-stage NSCLC patients with targeted therapy naïve (TN), on-treatment regressing or stable (residual disease, RD), or drug-resistant (progressive disease, PD) tumors was published by Maynard *et al*^24^ and obtained from https://github.com/czbiohub/scell_lung_adenocarcinoma. A Seurat object containing raw counts of filtered cells (S03_Merged_main_filtered_with_neo_osi.Rdata; nFeature_RNA >500 and nCount_RNA >50,000) was imported into R (v4.3.2)^63^ and further analyzed with Seurat (v5.0.3)^64^ after updating the Seurat object and its RNA assay to v5. Monocytes/macrophages (MoMacs) were identified by stepwise clustering of all cells, identified immune cells and identified mononuclear phagocytes (MNP). For dimensionality reduction and clustering, data was normalized by total counts per cell and log-transformed (NormalizeData, scale.factor = 10,000), searched for highly variable features (FindVariableFeatures), scaled (ScaleData, features = all) and subjected for PCA (RunPCA). FindNeighbors, FindClusters (resolution = 0.5, except for immune cells 0.7) and RunUMAP were run using first 20 PCs. Clusters were manually annotated based on established marker genes: Immune cells were identified as *PTPRC*^+^ clusters not expressing marker genes for fibroblasts (*DCN*, *COL1A1*), endothelial cells (*CD34*, *VWF*), melanocytes (*PMEL*) or epithelial cells (*EPCAM, KRT19*) (**Supplementary Figure S4A-C**). MNPs among immune cells were defined based on *CD68* expression after excluding neutrophil (*CXCR2, CSF3R*), mast cell (*MS4A2*, *TPSAB1*), pDC (*LILRA4*, *CLEC4C*), NKT (*CD3E*, *CD3D*, *NCAM1*) and B cell clusters (*CD79A*, *MS4A1*, *JCHAIN*) (Supplementary Figure S4D-F). After MNP subclustering, MoMacs were identified as *CD68*^+^*LYZ*^+^*S100A8*^+/low^ cells without expression of DC1 (*CLEC9A*^+^*XCR1*^+^*BTLA*^+^), DC2 (*CD1C*^+^*CLEC10A*^+^*FCER1A*^+^), LAMP3^+^ DC (*LAMP3*^+^*FSCN1*^+^*CCR7*^+^), Langerhans cells (*CD1A*^+^*CD207*^+^) or T cell doublet (*CD3E*^+^*CD3D*^+^) genes (Supplementary Figure S4G-I).^25,65,66^

Further analyses were performed for lung biopsy samples from patients with EGRF driver gene and at least 15 MoMacs per sample (n = 6 patients; n = 9 biopsies [3 TN, 5 RD, 1 PD]). All of these biopsies represented lung adenocarcinoma, and RD/PD patients were treated with osimertinib. MoMacs from these patients (n = 604 cells) were integrated by experimental batch (main cohort [n = 5, TH-] vs. additional neo-osi cohort [n = 4, AZ-]). Before integration, data was split by batch and subjected for normalization, feature identification, scaling and PCA as described above. After integration (IntegrateLayers, method = CCAIntegration), subclustering and UMAP dimensionality reduction were performed as described above (resolution = 0.8).

Differentially expressed genes (DEGs) in each MoMac cluster were identified by Wilcoxon rank sum test (FindAllMarkers, only.post = T, min.pct = 0.1) and upregulated DEGs were defined as genes with Padj <0.05 and log_2_FoldChange > 0.32. Pathway enrichment analysis for the obtained DEG lists was performed using clusterProfiler (v4.10.1)^67^ function compareCluster (fun = enrichGO) and GO biological process ontologies (org.Hs.eg.db, v3.18.0). Pathways passing adjusted p-value and q-value cut-offs (<0.05) were considered significant, and top 10 pathways were selected based on adjusted p-values. DEGs were re-analyzed with balanced TN and RD MoMac numbers per cluster in order to evaluate whether differences in patient MoMac numbers or the single PD sample affect MoMac cluster DEGs. For the balanced analysis, MoMacs from each TN and RD sample was randomly subseted to 10 (RD) or 15 (TN) MoMacs per cluster. The overlap between original and balanced analysis DEGs was visualized as Venn diagrams using package VennDiagram (v1.7.3).

Lung biopsy MoMacs were mapped to a cross-tissue single-cell atlas of human monocytes and macrophages, MoMac-VERSE^25^, using Seurat’s reference mapping functions: FindTransferAnchors (dims = 1:100, npcs = 100, normalization.method = “LogNormalize”) and MapQuery. MoMac-VERSE with UMAP reduction.model was kindly provided by Mulder and colleagues, while otherwise MoMac-VERSE is the same as found at https://github.com/gustaveroussy/FG-Lab. Gene set enrichment analysis was performed with escape (v2.0.0 and R v4.4.0)^68^ using Hallmark gene sets and ssGSEA algorithm. Resulting enrichment scores were normalized in each cell to number of uniquely expressed genes (performNormalization, scale.factor = nFeature_RNA, make.positive = T) and significance between TN and RD cells compared using Seurat’s Wilcoxon rank sum test. As normalization influenced enrichment scores between monocyte and macrophage clusters, we included only significant gene sets (adjusted P-value <0.05), where a similar difference was observed with unnormalized enrichment scores. Enrichment scores for these gene sets were plotted as a heatmap (heatmapEnrichment, scale = T, using normalized scores) and ridgeplots (ridgeEnrichment, scale = F, using normalized or unnormalized scores).

### CIBERSORTx immune cell deconvolution

Bulk RNA-seq expression data (FPKM) from EGFR-mutant lung cancer patient samples (GEO accession: GSE165019) were analyzed using CIBERSORTx^36^ with the LM22 immune cell signature matrix. Input files were formatted according to CIBERSORTx requirements, and all matched pre-and on-treatment samples provided in the dataset were included. Deconvolution was performed in absolute mode, with B-mode batch correction, no quantile normalization, and 100 permutations, using default settings for all other parameters. Estimated cell-type abundances for macrophages and T cells were extracted from the CIBERSORTx output, and correlations between inferred populations were calculated using Pearson correlation.

### Statistical analyses

Details of statistical analyses are provided in the figure legends. All tests were two-sided, and P < 0.05 was considered significant.

### Declaration of generative AI and AI-assisted technologies in the writing process

During the preparation of this work the authors used Grammarly and ChatGPT (GPT-5.1, OpenAI) in order to proofread and refine text for clarity. After using these tools, the authors reviewed and edited the content and take full responsibility for the content of the published article.

## Data accessibility

Previously published datasets re-analyzed in this study are publicly available and are cited within the manuscript. All data supporting the findings of this study are available from the corresponding author upon reasonable request.

## Acknowledgements

The authors wish to thank Orion Pharma for providing ORM-47286 for our experiments. We thank Maria Tuominen for skilled technical assistance. We thank the Turku Center for Disease Modelling (TCDM) for animal handling and the Histology core facility of the Institute of Biomedicine, University of Turku, Finland for histological stainings. Additionally, we want to thank the Proteomics core of Turku Bioscience Center for proteomics analyses, the Finnish Functional Genomics Centre supported by University of Turku, Åbo Akademi University, and Biocenter Finland for RNA sequencing services, CSC – IT Center for Science, Finland, for computational resources, the Genome Editing Core at the Turku Bioscience Center for providing assistance in CRISPR-Cas9 editing. This research was supported by funding from Finnish Cultural Foundation (K.J.K), Research Council of Finland (K.J.K; grant numbers 346656, 370023), Sigrid Jusélius Foundation (K.J.K), Instrumentarium Science Foundation (K.J.K), Jane and Aatos Erkko Foundation (K.J.K), Cancer Foundation Finland (K.J.K), Emil Aaltonen Foundation (M.H.), and iCANDOC (The Finnish National Doctoral Education Pilot in Precision Cancer Medicine) at University of Turku (J.B.).

## References

1. Boumahdi, S. & de Sauvage, F. J. The great escape: tumour cell plasticity in resistance to targeted therapy. Nature Reviews Drug Discovery 2019 19:1 19, 39–56 (2019).

2. Shen, S., Vagner, S. & Robert, C. Persistent Cancer Cells: The Deadly Survivors. Cell 183, 860–874 (2020).

3. Favaretto, G. et al. Neutrophil-activating secretome characterizes palbociclib-induced senescence of breast cancer cells. Cancer Immunol Immunother 73, 113 (2024).

4. Fleury, H. et al. Exploiting interconnected synthetic lethal interactions between PARP inhibition and cancer cell reversible senescence. Nature Communications 2019 10:1 10, 1–15 (2019).

5. Grimm, J. et al. BRAF inhibition causes resilience of melanoma cell lines by inducing the secretion of FGF1. Oncogenesis 7, 71 (2018).

6. Momeny, M. et al. DUSP6 inhibition overcomes neuregulin/HER3-driven therapy tolerance in HER2+ breast cancer. EMBO Mol Med 16, 1603–1629 (2024).

7. Kurppa, K. J. et al. Treatment-Induced Tumor Dormancy through YAP-Mediated Transcriptional Reprogramming of the Apoptotic Pathway. Cancer Cell 37, 104–122.e12 (2020).

8. Hagenbeek, T. J. et al. An allosteric pan-TEAD inhibitor blocks oncogenic YAP/TAZ signaling and overcomes KRAS G12C inhibitor resistance. Nature Cancer 2023 4:6 4, 812–828 (2023).

9. Chapeau, E. A. et al. Direct and selective pharmacological disruption of the YAP–TEAD interface by IAG933 inhibits Hippo-dependent and RAS–MAPK-altered cancers. Nature Cancer 2024 5:7 5, 1102–1120 (2024).

10. Zanconato, F., Cordenonsi, M. & Piccolo, S. YAP and TAZ: a signalling hub of the tumour microenvironment. Nature Reviews Cancer vol. 19 454–464 Preprint at 10.1038/s41568-019-0168-y (2019).

11. Guo, X. et al. Single tumor-initiating cells evade immune clearance by recruiting type II macrophages. Genes Dev 31, 247–259 (2017).

12. Ren, Z., Xu, Z., Chang, X., Liu, J. & Xiao, W. STC1 competitively binding βPIX enhances melanoma progression via YAP nuclear translocation and M2 macrophage recruitment through the YAP/CCL2/VEGFA/AKT feedback loop. Pharmacol Res 204, 107218 (2024).

13. Zhou, T. yi et al. Interleukin-6 induced by YAP in hepatocellular carcinoma cells recruits tumor-associated macrophages. J Pharmacol Sci 138, 89–95 (2018).

14. Thomann, S. et al. YAP-induced Ccl2 expression is associated with a switch in hepatic macrophage identity and vascular remodelling in liver cancer. Liver Int 41, 3011–3023 (2021).

15. Kim, W. et al. Hepatic Hippo signaling inhibits protumoural microenvironment to suppress hepatocellular carcinoma. Gut 67, 1692–1703 (2018).

16. Kleczko, E. K. et al. Novel EGFR-mutant mouse models of lung adenocarcinoma reveal adaptive immunity requirement for durable osimertinib response. Cancer Lett 556, 216062 (2023).

17. Dong, Z. Y. et al. EGFR mutation correlates with uninflamed phenotype and weak immunogenicity, causing impaired response to PD-1 blockade in non-small cell lung cancer. Oncoimmunology 6, e1356145 (2017).

18. Ercan, D. et al. Reactivation of ERK signaling causes resistance to EGFR Kinase inhibitors. Cancer Discov 2, 934–947 (2012).

19. Tricker, E. M. et al. Combined EGFR/MEK inhibition prevents the emergence of resistance in EGFR-mutant lung cancer. Cancer Discov 5, 960–971 (2015).

20. Milacic, M. et al. The Reactome Pathway Knowledgebase 2024. Nucleic Acids Res 52, D672–D678 (2024).

21. Martinez, F. O., Gordon, S., Locati, M. & Mantovani, A. Transcriptional Profiling of the Human Monocyte-to-Macrophage Differentiation and Polarization: New Molecules and Patterns of Gene Expression. The Journal of Immunology 177, 7303–7311 (2006).

22. Cassetta, L. & Pollard, J. W. Targeting macrophages: therapeutic approaches in cancer. Nature Reviews Drug Discovery 2018 17:12 17, 887–904 (2018).

23. Tap, W. D. et al. Structure-Guided Blockade of CSF1R Kinase in Tenosynovial Giant-Cell Tumor. New England Journal of Medicine 373, 428–437 (2015).

24. Maynard, A. et al. Therapy-Induced Evolution of Human Lung Cancer Revealed by Single-Cell RNA Sequencing. Cell 182, 1232–1251.e22 (2020).

25. Mulder, K. et al. Cross-tissue single-cell landscape of human monocytes and macrophages in health and disease. Immunity 54, 1883–1900.e5 (2021).

26. Nalio Ramos, R., et al. Tissue-resident FOLR2+ macrophages associate with CD8+ T cell infiltration in human breast cancer. Cell 185, 1189–1207.e25 (2022).

27. Timperi, E. et al. Lipid-Associated Macrophages Are Induced by Cancer-Associated Fibroblasts and Mediate Immune Suppression in Breast Cancer. Cancer Res 82, 3291–3306 (2022).

28. Metge, B. J. et al. Ribosomal RNA Biosynthesis Functionally Programs Tumor-Associated Macrophages to Support Breast Cancer Progression. Cancer Res 85, 1459–1478 (2025).

29. Caronni, N., La Terza, F., Frosio, L. & Ostuni, R. IL-1β+ macrophages and the control of pathogenic inflammation in cancer. Trends Immunol 46, 403–415 (2025).

30. Jaitin, D. A. et al. Lipid-Associated Macrophages Control Metabolic Homeostasis in a Trem2-Dependent Manner. Cell 178, 686–698.e14 (2019).

31. Patterson, M. T. et al. Trem2 promotes foamy macrophage lipid uptake and survival in atherosclerosis. Nature Cardiovascular Research 2023 2:11 2, 1015–1031 (2023).

32. Wang, Y. et al. TREM2-Mediated Cholesterol Efflux in Macrophages Inhibits Anti-Tumor Immunity via Limitation of CD4+ T and NK Cells. Advanced Science e06995 (2025) doi:10.1002/ADVS.202506995.

33. Shang, Y. et al. The transcriptional repressor Hes1 attenuates inflammation by regulating transcription elongation. Nat Immunol 17, 930–937 (2016).

34. Kim, M. S. et al. Disrupting Notch signaling related HES1 in myeloid cells reinvigorates antitumor T cell responses. Exp Hematol Oncol 13, 122- (2024).

35. Gurule, N. J. et al. A tyrosine kinase inhibitor-induced interferon response positively associates with clinical response in EGFR-mutant lung cancer. npj Precision Oncology 2021 5:1 5, 1–11 (2021).

36. Newman, A. M. et al. Determining cell type abundance and expression from bulk tissues with digital cytometry. Nature Biotechnology 2019 37:7 37, 773–782 (2019).

37. Lin, L. et al. The Hippo effector YAP promotes resistance to RAF- and MEK-targeted cancer therapies. Nature Genetics 2015 47:3 47, 250–256 (2015).

38. Shi, J. et al. The HER4-YAP1 axis promotes trastuzumab resistance in HER2-positive gastric cancer by inducing epithelial and mesenchymal transition. Oncogene 2018 37:22 37, 3022–3038 (2018).

39. Yun, M. R. et al. Targeting YAP to overcome acquired resistance to ALK inhibitors in ALK-rearranged lung cancer. EMBO Mol Med 11, e10581 (2019).

40. Nam, A. R. et al. YAP as a therapeutic target to reverse trastuzumab resistance. Gastric Cancer 1–15 (2025) doi:10.1007/S10120-025-01630-W/FIGURES/5.

41. Haderk, F. et al. Focal adhesion kinase-YAP signaling axis drives drug-tolerant persister cells and residual disease in lung cancer. Nat Commun 15, (2024).

42. Uhlén, M. et al. Tissue-based map of the human proteome. Science (1979) 347, (2015).

43. Ahlmark, M. et al. Tead inhibitors. (2025). EPO patent EP4323338B1, October 8, 2025.

44. Hata, A. N. et al. Tumor cells can follow distinct evolutionary paths to become resistant to epidermal growth factor receptor inhibition. Nat Med 22, 262–269 (2016).

45. Lee, C. K. et al. Checkpoint Inhibitors in Metastatic EGFR-Mutated Non–Small Cell Lung Cancer—A Meta-Analysis. Journal of Thoracic Oncology 12, 403–407 (2017).

46. Isomoto, K. et al. Impact of EGFR-TKI treatment on the tumor immune microenvironment in EGFR mutation-positive non-small cell lung cancer. Clinical Cancer Research 26, 2037–2046 (2020).

47. Han, R. et al. Tumour microenvironment changes after osimertinib treatment resistance in non-small cell lung cancer. Eur J Cancer 189, (2023).

48. Gainor, J. F. et al. EGFR mutations and ALK rearrangements are associated with low response rates to PD-1 pathway blockade in non-small cell lung cancer: A retrospective analysis. Clinical Cancer Research 22, 4585–4593 (2016).

49. Fang, Y. et al. Comprehensive analyses reveal TKI-induced remodeling of the tumor immune microenvironment in EGFR/ALK-positive non-small-cell lung cancer. Oncoimmunology 10, (2021).

50. Rannikko, J. H. & Hollmén, M. Clinical landscape of macrophage-reprogramming cancer immunotherapies. British Journal of Cancer 2024 1–14 (2024) doi:10.1038/s41416-024-02715-6.

51. Xu, J. et al. Dual roles and therapeutic targeting of tumor-associated macrophages in tumor microenvironments. Signal Transduction and Targeted Therapy 2025 10:1 10, 268- (2025).

52. Kim, S. et al. Remodeling of tumor microenvironments by EGFR tyrosine kinase inhibitors in EGFR-mutant non-small cell lung cancer. iScience 28, 111736 (2025).

53. Zhang, B. et al. M2-polarized macrophages contribute to the decreased sensitivity of EGFR-TKIs treatment in patients with advanced lung adenocarcinoma. Medical Oncology 31, 127-(2014).

54. Okawa, S. et al. Colony-Stimulating Factor-1 Receptor Inhibitor Augments Osimertinib-Induced Antitumor Immunity via Suppression of Macrophages in Lung Cancer Harboring EGFR Mutation. Mol Cancer Ther 24, 1763–1774 (2025).

55. Du, X. X. et al. YAP/STAT3 promotes the immune escape of larynx carcinoma by activating VEGFR1-TGFβ signaling to facilitate PD-L1 expression in M2-like TAMs. Exp Cell Res 405, 112655 (2021).

56. Meli, V. S. et al. YAP-mediated mechanotransduction tunes the macrophage inflammatory response. Sci Adv 6, (2020).

57. Sapudom, J., Alatoom, A., Tipay, P. S. & Teo, J. C. Matrix stiffening from collagen fibril density and alignment modulates YAP-mediated T-cell immune suppression. Biomaterials 315, 122900 (2025).

58. Wang, G. et al. Targeting YAP-dependent MDSC infiltration impairs tumor progression. Cancer Discov 6, 80–95 (2016).

59. Bankhead, P. et al. QuPath: Open source software for digital pathology image analysis. Scientific Reports 2017 7:1 7, 1–7 (2017).

60. Love, M. I., Huber, W. & Anders, S. Moderated estimation of fold change and dispersion for RNA-seq data with DESeq2. Genome Biol 15, (2014).

61. Frankenfield, A. M., Ni, J., Ahmed, M. & Hao, L. Protein Contaminants Matter: Building Universal Protein Contaminant Libraries for DDA and DIA Proteomics. J Proteome Res 21, 2104–2113 (2022).

62. Callister, S. J. et al. Normalization approaches for removing systematic biases associated with mass spectrometry and label-free proteomics. J Proteome Res 5, 277–286 (2006).

63. R Core Team. R: A Language and Environment for Statistical Computing.

64. Hao, Y. et al. Dictionary learning for integrative, multimodal and scalable single-cell analysis. Nat Biotechnol 42, 293–304 (2024).

65. Cheng, S. et al. A pan-cancer single-cell transcriptional atlas of tumor infiltrating myeloid cells. Cell 184, 792–809.e23 (2021).

66. Bassez, A. et al. A single-cell map of intratumoral changes during anti-PD1 treatment of patients with breast cancer. Nat Med 27, 820–832 (2021).

67. Wu, T. et al. clusterProfiler 4.0: A universal enrichment tool for interpreting omics data. Innovation (Cambridge (Mass.)) 2, (2021).

68. Borcherding, N., et al. Mapping the immune environment in clear cell renal carcinoma by single-cell genomics. Commun Biol 4, (2021).

