## Supplementary Figures for "YAP/TEAD drives treatment-induced adaptive immunosuppression in EGFR-mutant lung cancer"

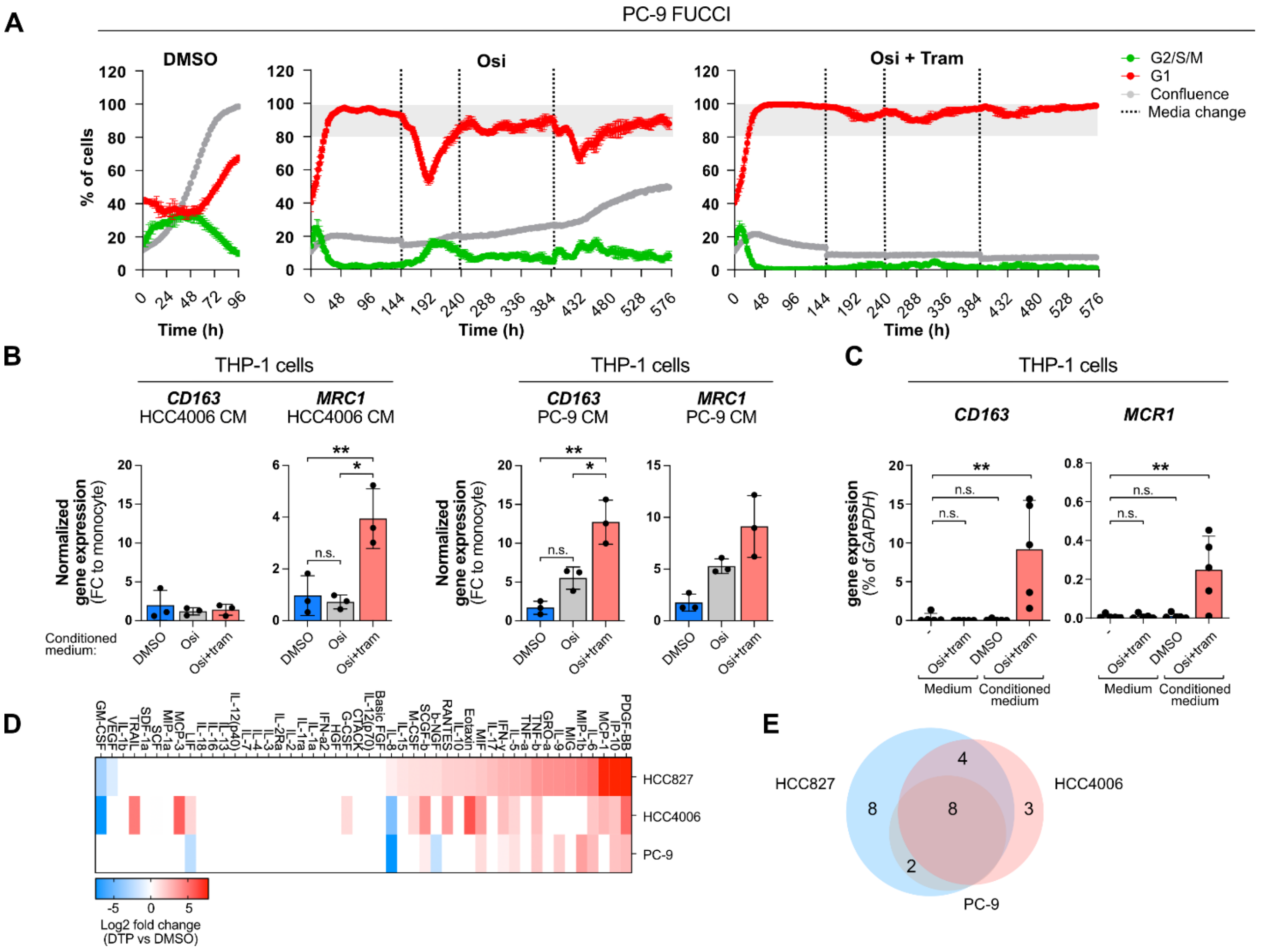

**Supplementary Figure 1. Secreted factors from quiescent, on-treatment cancer cells reprogram macrophages to an M2-like immunosuppressive state.** A) PC-9 cells expressing pLEX307-FUCCI cell cycle reporter were treated with either DMSO, 100 nM osimertinib alone, or in combination with 30 nM trametinib for 24 days. B) THP-1 cells were treated for 6 days with conditioned media collected from HCC4006 and PC9 cells treated either with DMSO (48 hours), 100 nM osimertinib alone (O), or in combination with 30 nM trametinib (O+T for 12 days), and the expression of M2 marker genes was analyzed with RT-qPCR. C) THP-1 cells were treated as in (B) with either HCC827-conditioned media or RPMI-1940 ± O+T. D) Immunomodulatory secretome in HCC827, HCC4006, and PC9 cells analysed with Bio-Plex Pro Human Cytokine 27-plex assay. Quantification of secreted proteins is shown as log2 fold change in DTP vs DMSO cells. E) Overlap of secreted proteins in all three cell lines from (D). One-way ANOVA was used to assess statistical significance. **, P-value <0.01; *, P-value < 0.05

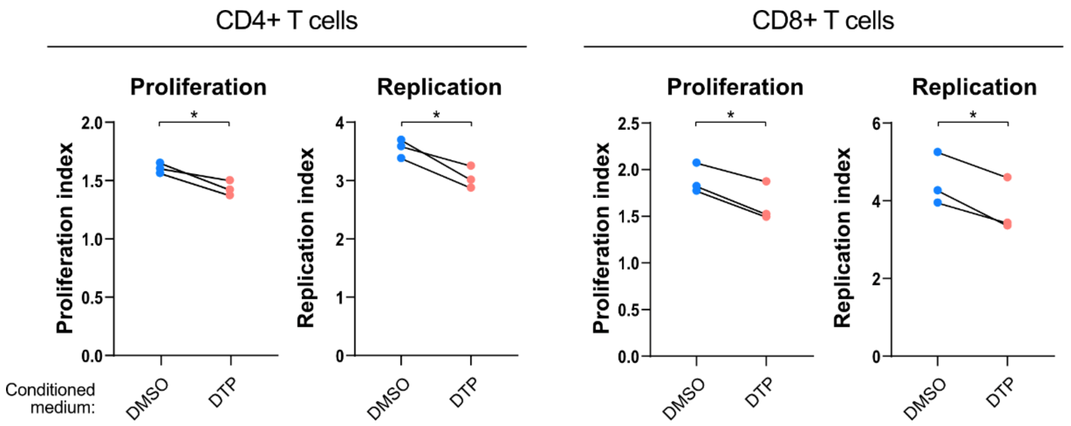


**Supplementary Figure 2. Proliferation and replication indices of CD4⁺ and CD8⁺ T cells co-cultured with DTP-reprogrammed monocytes.** Proliferation and replication indices were calculated from flow cytometry division tracking data corresponding to Figure 3B–C. Monocytes were exposed to DTP or control conditioned medium for 6 days prior to co-culture with pre-activated, donor-matched $CD4^+$ and $CD8^+$ T cells for 72 hours. A t-test was used to assess statistical significance. DTP = HCC827 cells treated with 100 nM osimertinib + 30 nM trametinib for 12 days.

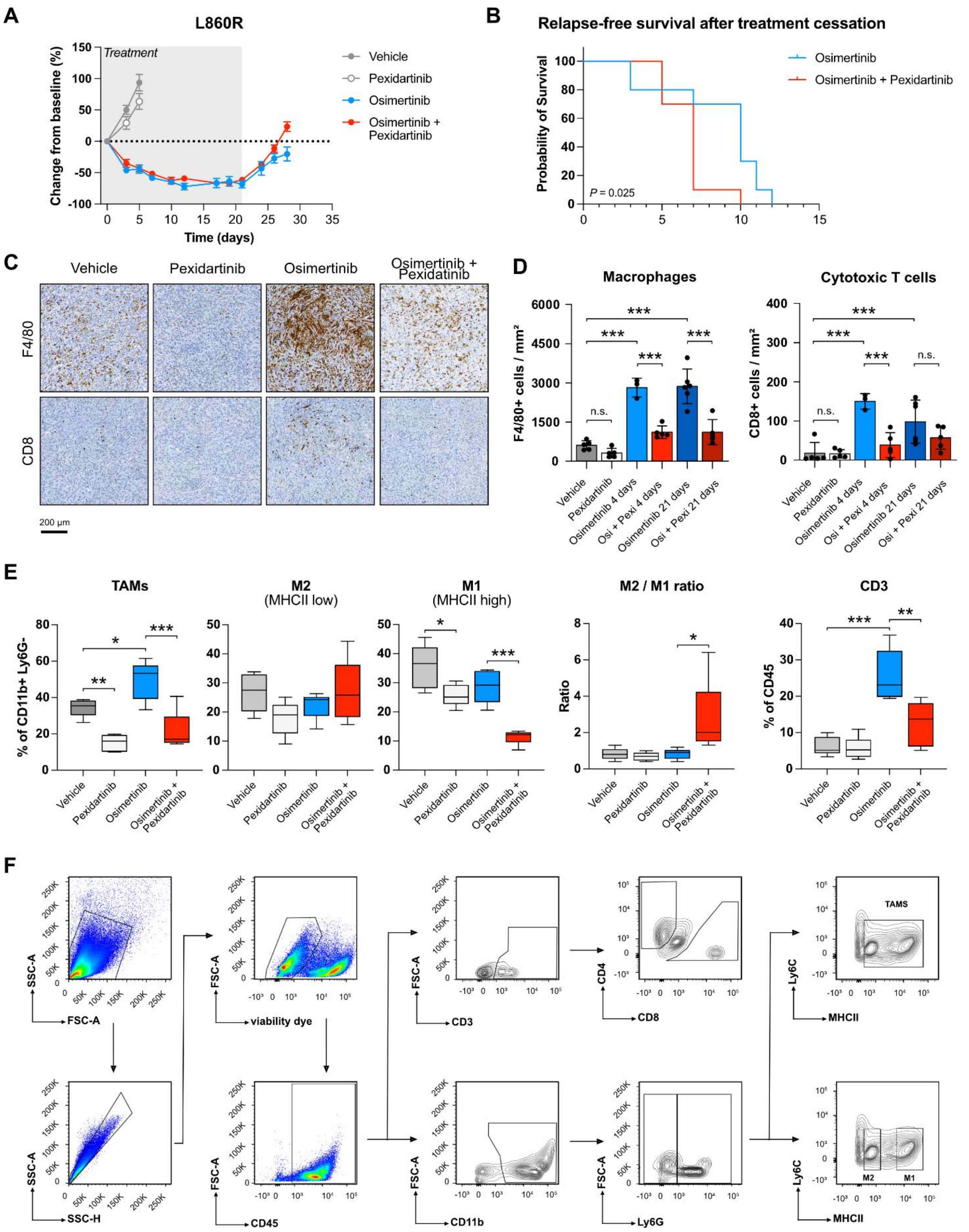


**Supplementary Figure 3. CSF1R inhibition does not enhance osimertinib efficacy in EGFR-mutant syngeneic mouse model.** A) Osimertinib ± pexidartinib response in syngeneic EGFR-mutant lung cancer model L860R in C57Bl/6 mice. B) Relapse-free survival of treated mice, determined as time to tumor regrowth after treatment cessation to the original size at day 0. C) Immunohistochemical staining of tumor samples from (A). F4/80, macrophages; CD8a, cytotoxic T-cells. D) Quantification of positive cells detected per mm2 in tumors depicted in (C) using QuPath. E) Flow cytometry analysis of tumours after 4 days of treatment. F) Flow cytometry density plots showing the comprehensive gating strategy used to detect different immune cell populations in mouse tumors. Data collected was analysed with FlowJo software. One-way ANOVA was used to assess statistical significance (C, D). ***, P-value <0.001; **, P-value <0.01; *, P-value < 0.05

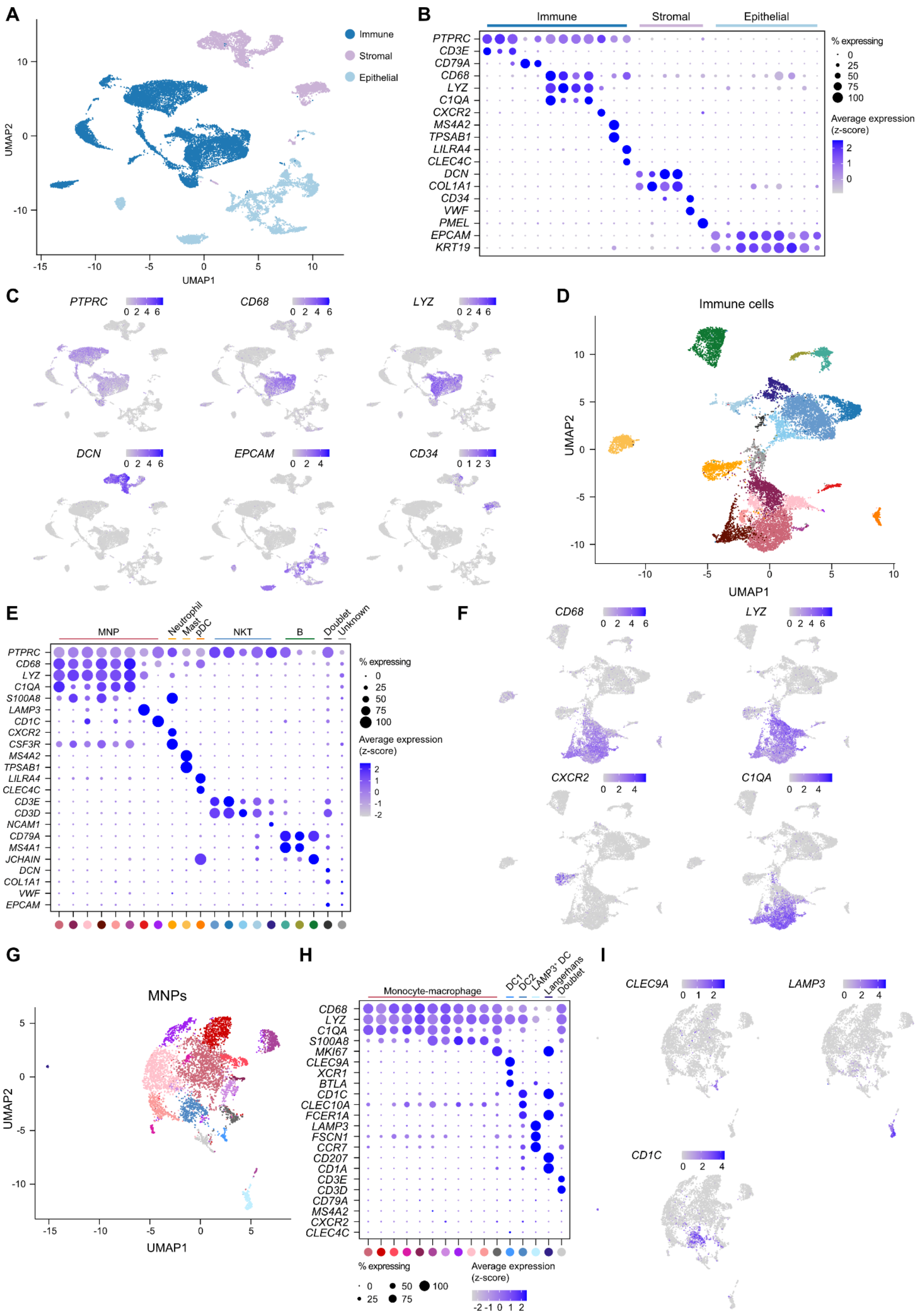

A
Immune
Stromal
Epithelial
UMAP1
UMAP2
B
PTPRC
CD3E
CD79A
CD68
LYZ
C1QA
CXCR2
MS4A2
TPSAB1
LILRA4
CLEC4C
DCN
COL1A1
CD34
VWF
PMEL
EPCAM
KRT19
% expressing
Average expression (z-score)
C
D
Immune cells
E
MNP
Neutrophil
Mast
pDC
NKT
B
Doublet
Unknown
S100A8
LAMP3
CD1C
CSF3R
CD3D
NCAM1
MS4A1
JCHAIN
F
G
MNPs
H
Monocyte-macrophage
DC1
DC2
LAMP3⁺ DC
Langerhans
Doublet
MKI67
CLEC9A
XCR1
BTLA
CLEC10A
FCER1A
FSCN1
CCR7
CD207
CD1A
I

**Supplementary Figure 4. Clustering of scRNA-sequenced cells from metastatic NSCLC biopsies to identify monocytes and macrophages, related to Figure 4.** A) UMAP plot showing the clustering of all NSCLC biopsy cells passing initial quality filters (n = 23,479 cells), colored by main cell type (immune, stromal or epithelial). B) A dot plot displaying average cluster marker expression in immune, stromal and epithelial cell clusters from A. C) UMAP plots colored by the indicated immune (*PTPRC*, *CD68, LYZ*), stromal (*DCN, CD34*) or epithelial (*EPCAM*) marker gene expression. D) Subclustering of immune cells (n = 13,611 cells) identified in A-C. E) Average expression of immune cell marker genes in clusters from D. Mononuclear phagocytes were identified as positive for monocyte/macrophage (*CD68, LYZ, C1QA, S100A8*) or dendritic cell (*CD1C, LAMP3*) marker genes. F) UMAP plot from D colored by mononuclear phagocyte (*CD68*, *LYZ*, *C1QA*) or neutrophil (*CXCR2*) marker genes. G) UMAP plot showing the subclustering of mononuclear phagocytes (n = 5,066 cells) identified in D-F. H) Average expression of monocyte, macrophage and dendritic cell marker genes in MNP clusters from G. After excluding the indicated clusters of dendritic cells, Langerhans cells and T cell doublets, remaining $CD68^{+}LYZ^{+}C1QA^{+/-}S100A8^{+/-}$ clusters were identified as monocytes/macrophages. I) UMAP plot from G colored by dendritic cell marker gene expression. B = B cell; (p)DC = (plasmacytoid) dendritic cell; MNP = mononuclear phagocyte; NKT = NK or T cell; UMAP = uniform manifold approximation and projection.

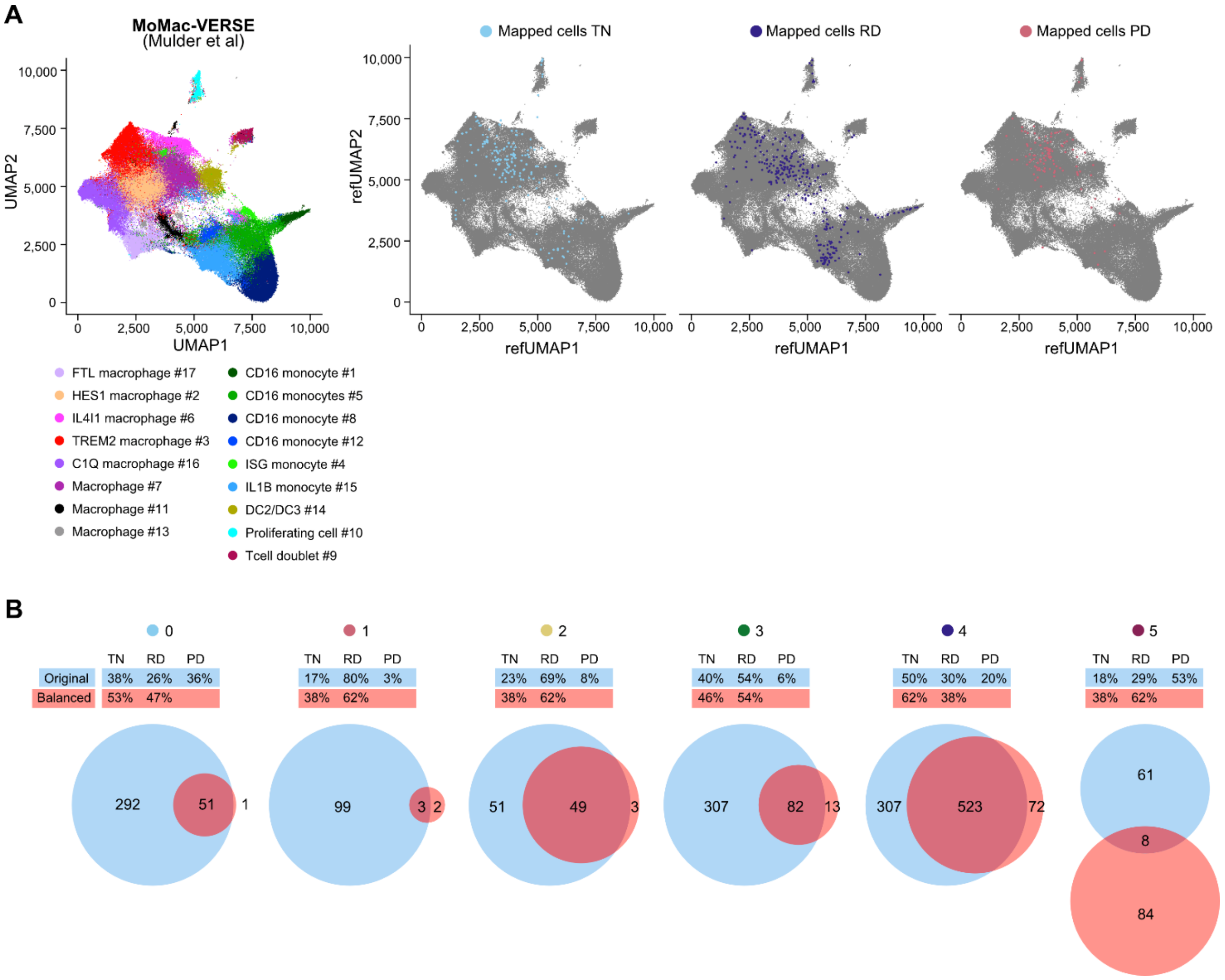


**Supplementary Figure 5.** A) Original MoMac-VERSE UMAP plot (left) and mapped cells from TN, RD and PD samples highlighted on top of the original MoMac-VERSE UMAP colored in gray (right). Related to Figure 4E-F. B) MoMac cluster DEGs were re-analyzed after excluding PD sample and balancing TN and RD MoMac numbers in each cluster. Venn diagrams show DEG overlap between balanced re-analysis and original analysis presented in Figure 4D and Supplementary Table S1. Percentages show original and balanced analysis cell numbers in each cluster. PD = progressive disease; RD = residual disease; TN = targeted therapy naïve.

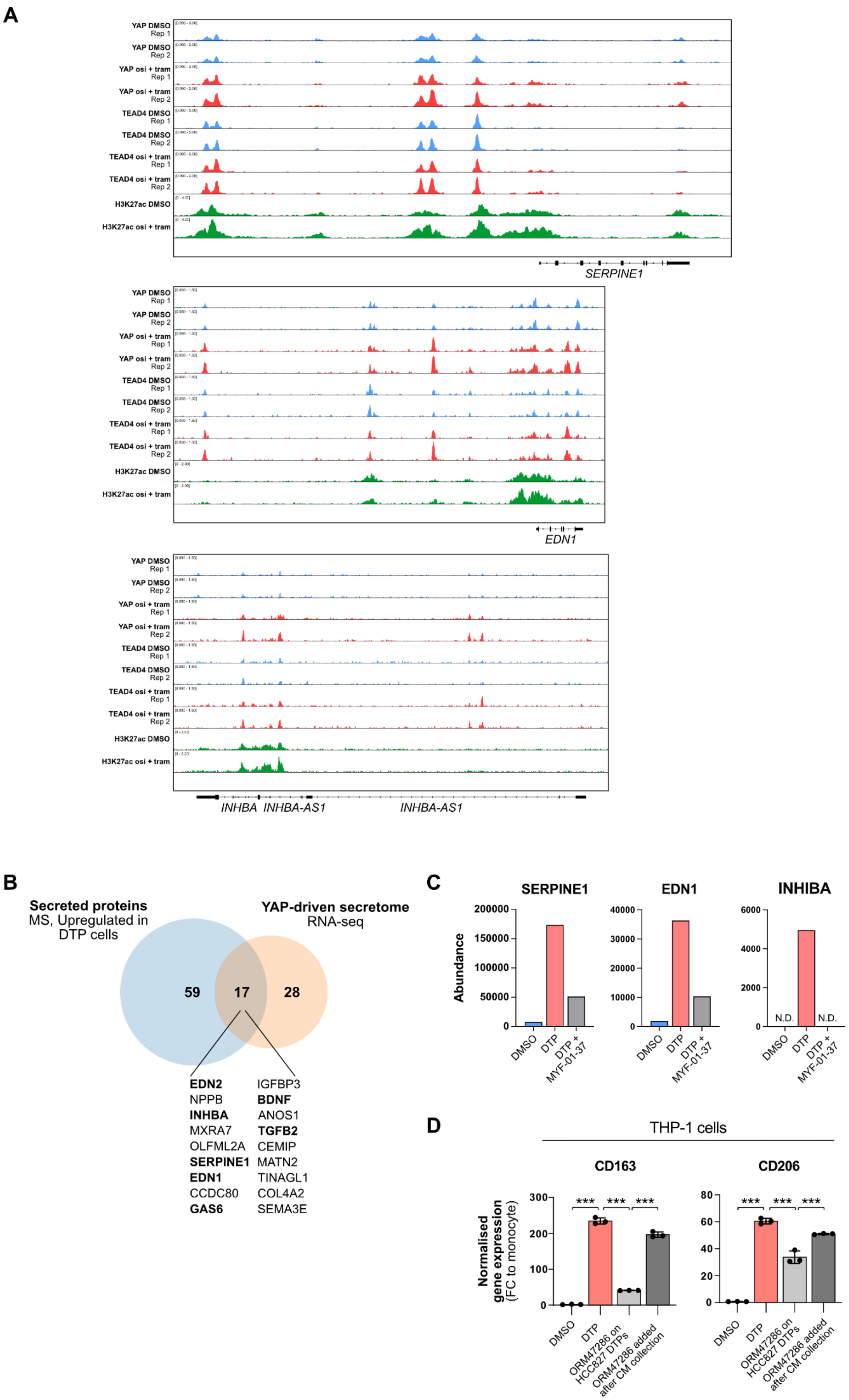
A
YAP DMSO Rep 1
YAP DMSO Rep 2
YAP osi + tram Rep 1
YAP osi + tram Rep 2
TEAD4 DMSO Rep 1
TEAD4 DMSO Rep 2
TEAD4 osi + tram Rep 1
TEAD4 osi + tram Rep 2
H3K27ac DMSO
H3K27ac osi + tram
SERPINE1
EDN1
INHBA
INHBA-AS1
INHBA-AS1
B
Secreted proteins
MS, Upregulated in DTP cells
YAP-driven secretome
RNA-seq
59
17
28
EDN2
NPPB
INHBA
MXRA7
OLFML2A
SERPINE1
EDN1
CCDC80
GAS6
IGFBP3
BDNF
ANOS1
TGFB2
CEMIP
MATN2
TINAGL1
COL4A2
SEMA3E
C
SERPINE1
EDN1
INHBA
Abundance
200000
150000
100000
50000
0
40000
30000
20000
10000
0
6000
4000
2000
0
N.D.
N.D.
DMSO
DTP
DTP + MYF-01-37
D
THP-1 cells
CD163
CD206
Normalised gene expression (FC to monocyte)
300
200
100
0
80
60
40
20
0
***
***
***
DMSO
DTP
ORM47286 on HCC827 DTPs
ORM47286 added after CM collection

**Supplementary Figure 6.** A) Representative images demonstrating YAP and TEAD binding to the loci of YAP/TEAD-driven secretome genes. B) Overlap of YAP/TEAD-driven secretome of HCC827 and HCC4006 DTPs from Figure 5D with DTP secretome (HCC827) from proteomics analysis. C) Abundance of selected proteins from (B) in HCC827 conditioned media. D) THP-1 cells were treated with HCC827 conditioned media as in Supplementary Figure 1B. HCC827 DTPs were treated with 1 μM ORM-47286, or 1 μM ORM-47286 was added to collected condition media of O+T treated HCC827 DTPs before exposure to THP-1 cells. One-way ANOVA was used to assess statistical significance (C). ***, P-value <0.001; **, P-value <0.01.
